# Construction of Disease-specific Cytokine Profiles by Associating Disease Genes with Immune Responses

**DOI:** 10.1101/2021.09.10.459816

**Authors:** Tianyun Liu, Shiyin Wang, Russ B Altman

**Affiliations:** Department of Bioengineering, Stanford University, Stanford, CA; Chinese Undergraduate Visiting Research Program, Stanford University, Stanford, CA

## Abstract

The pathogenesis of many inflammatory diseases is a coordinated process involving metabolic dysfunctions and immune response—usually modulated by the production of cytokines and associated inflammatory molecules. In this work, we seek to understand how genes involved in pathogenesis which are often not associated with the immune system in an obvious way communicate with the immune system. We have embedded a network of human protein-protein interactions (PPI) from the STRING database with 14,707 human genes using feature learning that captures high confidence edges. We have found that our predicted Association Scores derived from the features extracted from STRING’s high confidence edges are useful for predicting novel connections between genes, thus enabling the construction of a full map of predicted associations for all possible pairs between 14,707 human genes. In particular, we analyzed the pattern of associations for 126 cytokines and found that the six patterns of cytokine interaction with human genes are consistent with their functional classifications. In order to define the disease-specific roles of cytokines we have collected gene sets for 11,944 diseases from DisGeNET. We used these gene sets to predict disease-specific gene associations with cytokines by calculating the normalized average Association Scores between disease-associated gene sets and the 126 cytokines; this creates a unique profile of inflammatory genes (both known and predicted) for each disease. We validated our predicted cytokine associations by comparing them to known associations for 171 diseases. The predicted cytokine profiles correlate (p-value<0.05) with the known ones in 147 diseases. We further characterized the profiles of each disease by calculating an “Inflammation Score” that summarizes different modes of immune responses. Finally, by analyzing subnetworks formed between disease-specific pathogenesis genes, hormones, receptors, and cytokines, we identified the key genes responsible for interactions between pathogenesis and inflammatory responses. These genes and the corresponding cytokines used by different immune disorders suggest unique targets for drug discovery.

## Introduction

The pathogenesis of inflammatory diseases is a coordinated process involving metabolic dysfunctions, signaling, and innate immune response—modulated by the production of cytokines and associated inflammatory molecules. The continued discovery of novel pathways and inflammatory mediators reveals the complexity and richness of autoimmune diseases ^1 2^, but the complete molecular decision network behind these processes and the coordination between cytokine signaling and underlying disease biology are not understood ^3 4 5^. Many models posit a common cytokine framework with highly conserved mechanisms of inflammation^6,7^. Recent advances in genome-wide association studies (GWAS) provide evidence of considerable genetic overlap between autoimmune diseases, along with unique loci for individual diseases-- and sometimes their subtypes ^8 9, 10^. The success of anti-TNF treatment in multiple inflammatory diseases suggests that there is a shared cytokine framework (at least for a subset of diseases) that defines conserved mechanisms of inflammation ^11^. However, clinical trials testing the efficacy of cytokine inhibitors suggest a more complex set of interacting cytokine mechanisms that are associated with different disease phenotypes ^12^. For example, the Jak-Stat-Socs signaling module can have either pro- and antiinflammatory outcomes depending on the activation pattern of cytokine receptors—and different diseases show different patterns. If diseases do not have the same cytokine activity profile, then it is important to define which ones are key for each disease. Therefore, we ask the question: What is the degree to which cytokine responses are shared across diseases or specific to each disease? And how does heterogeneity of cytokine responses mediate different pathogenesis and inflammatory processes?

In order to address the above questions, we require a deeper understanding of the connections between cytokine signaling and the disease-specific genes implicated in pathogenesis of specific diseases and discovered through GWAS or transcriptional studies. Networks of interacting genes provide a useful representation of the functional associations between genes and gene modules ^13 14^. Unfortunately, efforts to identify disease-associated genes do not always provide a clear link between pathogenesis and immune response. Publicly available immunology databases often limit their data in the immune process. ImmuneSigDB identifies changes in expression of sets of genes corresponding to inflammation, enabling a system analysis of data to improve the understanding of immune processes ^15^. ImmProt^16^ applied high-resolution mass spectrometry-based proteomics to characterize primary human immune cell types for ligand and receptor interactions, thereby connecting distinct immune functions. ImmuneXpresso ^17^ identifies relationships between cells, cytokines and diseases via Natural Language Processing (NLP).

Ideally, we would like to map known immune response genes more completely to human gene networks to better identify potential links to pathogenesis genes. STRING is a comprehensive gene network that focuses chiefly on molecular pathways, with cancer heavily emphasized; unfortunately, STRING does not provide fully elaborated links to immune response^18^. We have compared the network sparsity (the ratio of the number of actual links to the total number of links present in a complete graph) within and between known immune and disease-related functional modules in STRING (Supplementary Material Figure S1). We have found that the associations between different immune response genes (as identified by ImmProt) are under-represented. Either the connections are more difficult to study and characterize or that the connections are less dense compared with metabolic or signaling modules as the human body requires more traffic for metabolic or signaling activities.

Nevertheless, we aim to bridge the gap between pathogenesis and the immune processes that ultimately cause systemic damage. This will allow us to understand how cytokine responses are triggered in a disease-specific manner. We hypothesize that specific clinical phenotypes result from the interactions between disease-specific cytokines and disease-related genes (identified through genetics, transcriptomics, and analysis of metabolic dysfunctions), even though they also may share a common cytokine elements and conserved mechanisms of inflammation. We identify these disease-specific cytokines and their associated disease-specific genes to provide insights into the underlying molecular mechanisms. These mechanisms may suggest new approaches to treatment and treatment combinations for specific clinical phenotypes.

## Results

### A complete map of associations between 14K human genes

We developed a novel network method to construct a complete interaction map between human genes by embedding a PPI of 14,707 human genes using network scalable feature learning ^19^ that captures 728,090 high confidence edges in STRING ^18^ (Figure 1A and Supplementary Material Table S1). We have calculated Association Scores of all possible pairs (108,140,571 pairs) between the 14,707 human genes. The distribution of the predicted Association Scores of all possible pairs (108,140,571 pairs) is similar with that of the known pairs in STRING (9,250,034 pairs) (Supplementary Material Figure S2-S4). Among these pairs, STRING provides a confidence index for 9,250,034 pairs, of which 8,521,944 have a confidence index below 800 (the STRING cutoff) and were not used for embedding. Figure 2 shows that the predicted Association Scores correlate with the level of confidence in STRING. The 9,250,034 pairs are grouped into four boxplots based on their predicted Association Scores (Figure 2A). As the predicted scores increase, the average STRING confidence index also increase. When the predicted scores are below 0.6, the STRING confidence is low with an average around 200. When the predicted scores are above 0.8, 76% of the pairs have confidence above 700, and 75% of the pairs above 800 confidence cutoffs (Figure 2B). Thus, it appears that the Association Scores between genes in an embedded network are good predictors of protein-protein interactions, and thus our embedding enables the construction of a complete and reliable network between 14,707 human genes.

**Figure 1.**
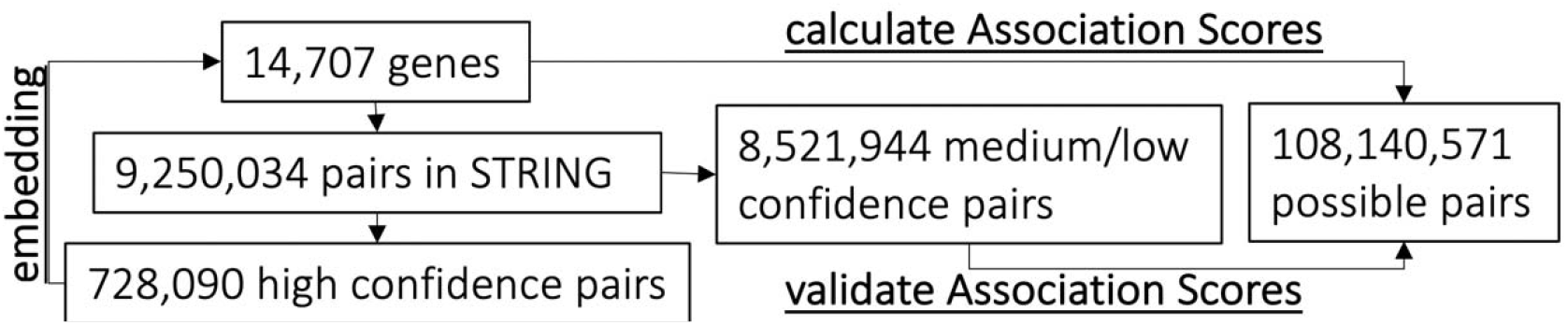
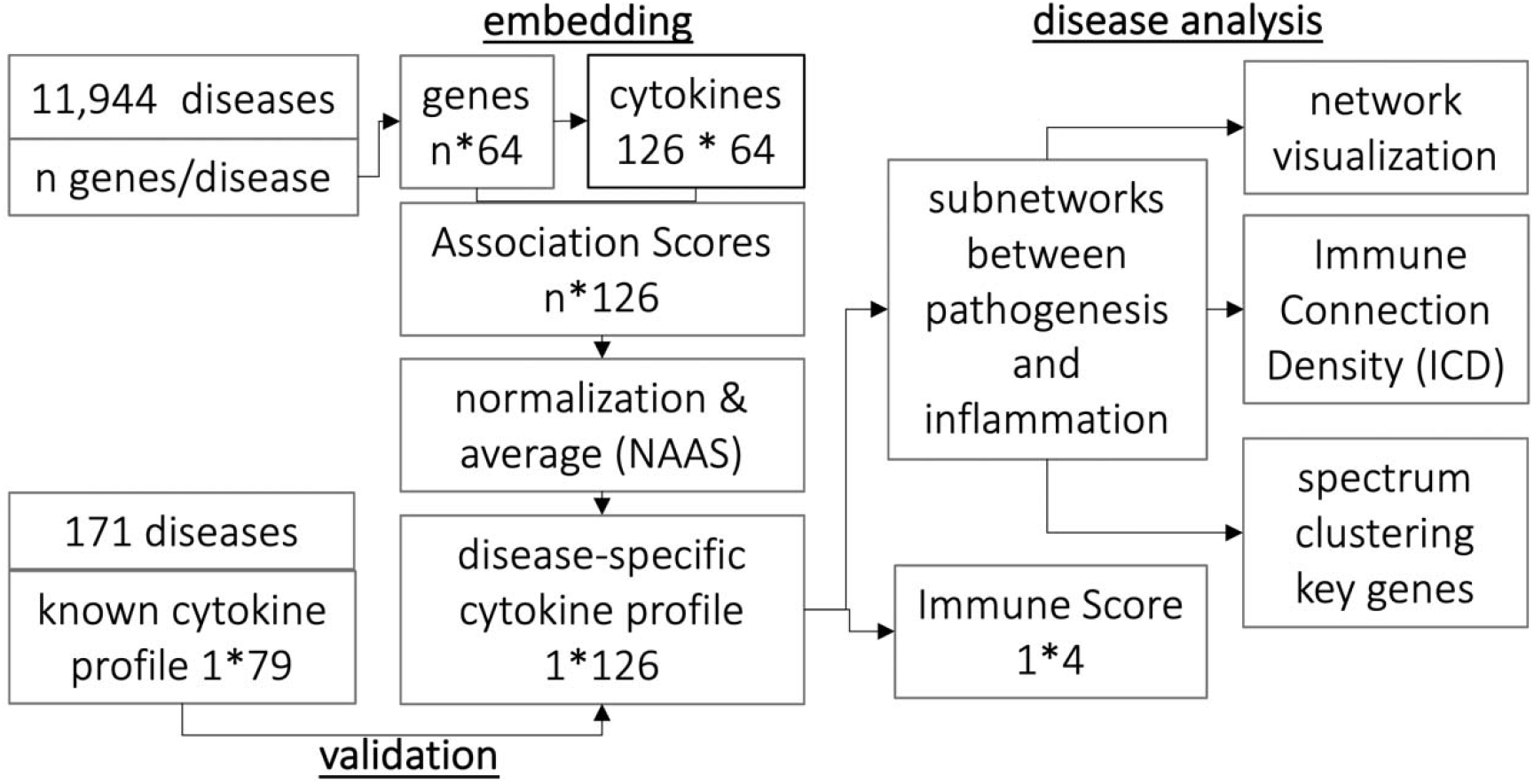
Data flow for (A) predicting Association Scores and (B) analyzing on disease-specific cytokine profiles. A. Network features of the high confidence STRING pairs were used to embed 14,707 human genes. The predicted associations between the 14,707 genes were validated by medium or low confidence STRING pairs. B. Given each gene associated with a given disease, we calculated Association Scores with the 126 cytokines. The Association Scores were averaged and normalized to NAAS that represent the cytokine profile of the given disease. The profiles were further analyzed by (1) calculating Immune Scores and (2) analyzing subnetworks formed between pathogenesis and inflammation by employing network visualization, spectrum partition, and estimation of connection density.

**Figure 2.**
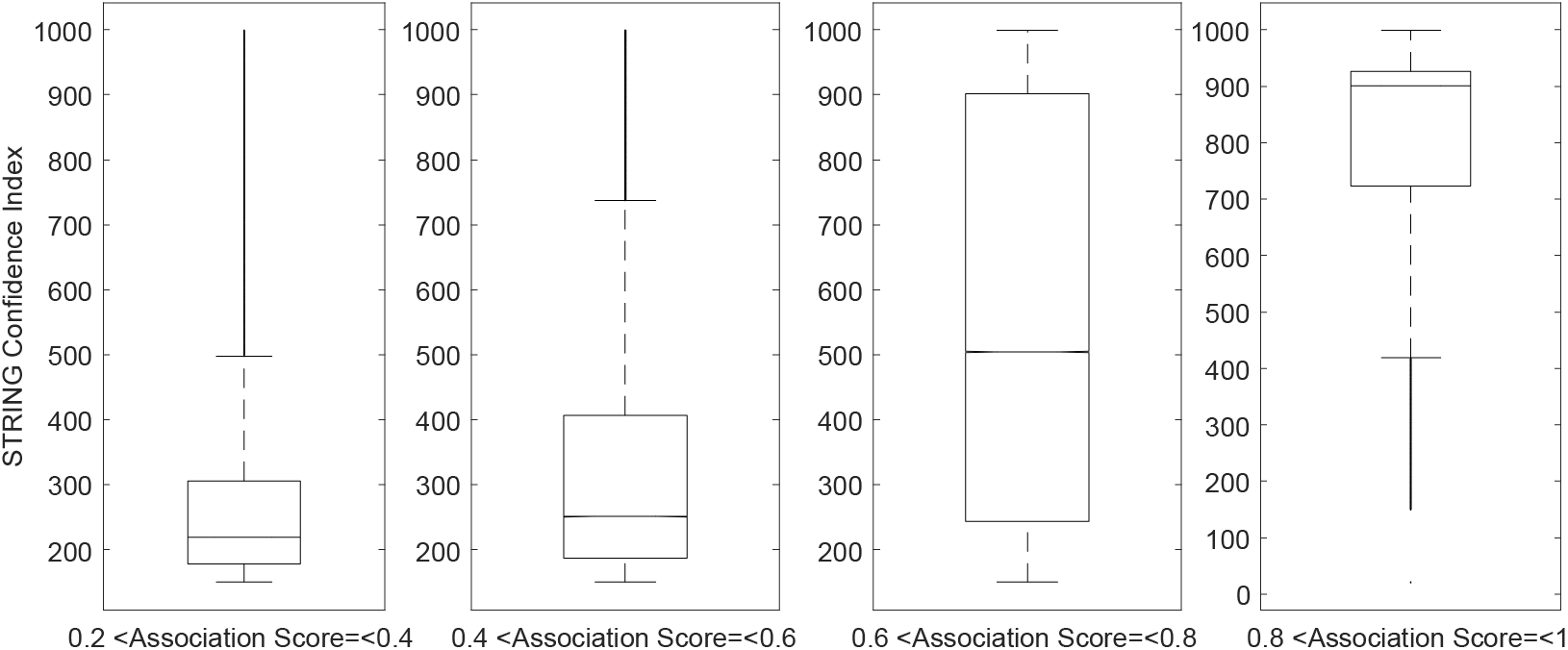
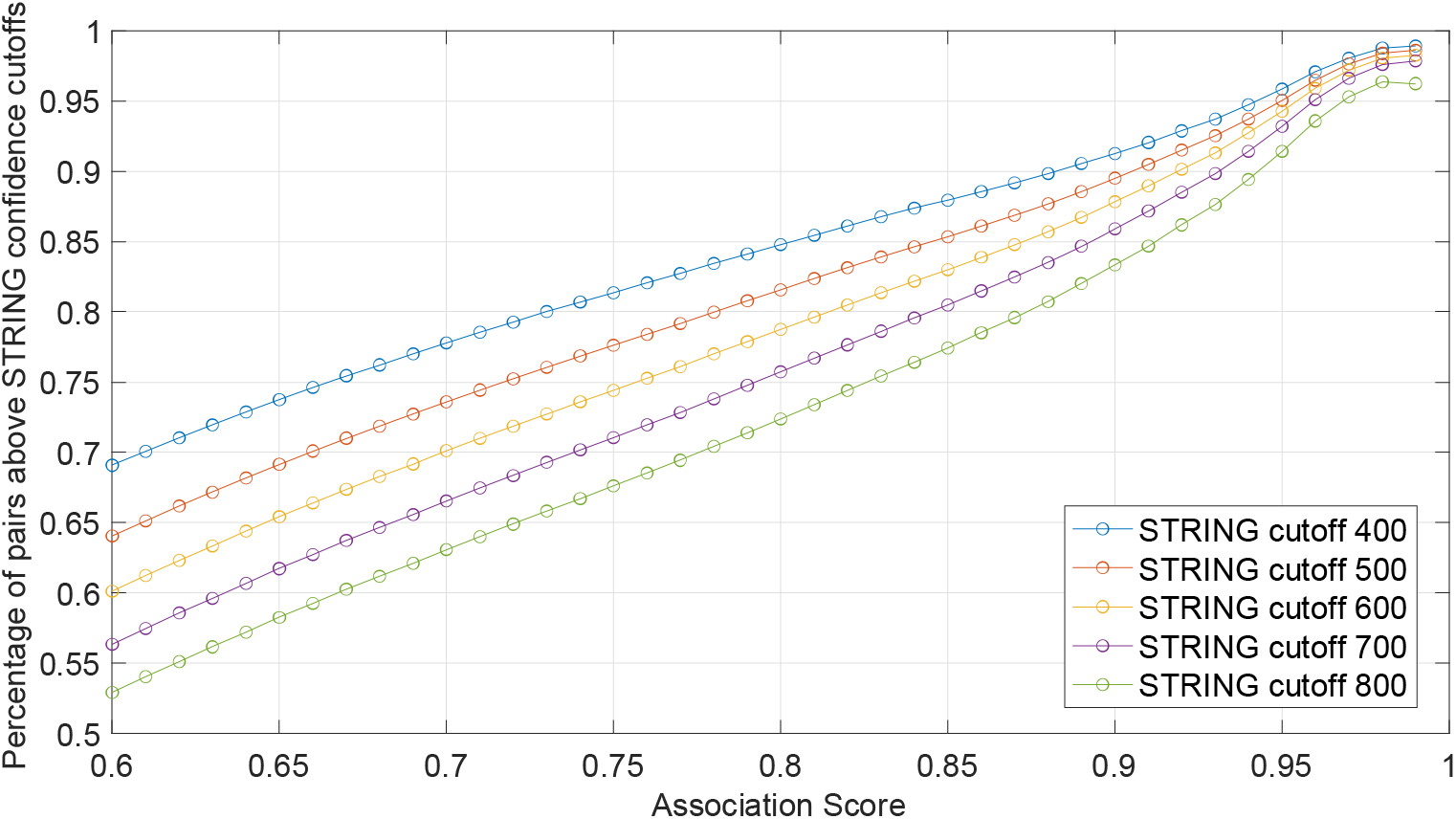
The predicted Association Score indicates the confidence of associations. We calculated the Association Scores for all possible pairs (108,140,571 pairs) between the 14,707 human genes. A total of 9,250,034 edges are known with confidence index (discrete scores) in STRING. A. These pairs are shown in four boxplots based on their predicted Association Scores. Along the increase of the Association Scores, the average STRING confidence index increase. B. The percentage of pairs above a certain confidence index cutoff (400 to 800).

### Essential cytokines can be classified into six clusters based on their interaction profiles

We identified 126 well-studied cytokines based on ImmuneXpresso ^17^ in our embedded network. We call these 126 cytokines “essential cytokines” because they are linked with at least one disease in the literature. The term “cytokine” encompasses inflammatory regulators, including interferons, the interleukins, the chemokine family, mesenchymal growth factors, the tumor necrosis factor family and other non-classified ones^20^. Each cytokine can be described by its location in the embedded network space, and their Association Scores to the 14,581 non-cytokine human genes suggest known and novel interactions with other human genes (Figure 1B). The Association Scores between each of the cytokines and the 14,581 non-cytokine human genes classified these 126 cytokines generally into six clusters, which we have named based on the types of the most enriched cytokine/chemokines: TGF-CLU (growth factors), Chemokine-CLU (chemokines), TNF-CLU (TNFs), IFN-CLU (interferons), IL-CLU (interleukins), and Unclassified-CLU (Figure 3). However, it remains difficult to quantify the enrichment and purity of these clusters, due to the pleiotropy and element of redundancy in the cytokine family (each cytokine has many overlapping functions, and with each function mediated by more than one cytokine.) Six sets of close interactors are suggested by the dendrogram of the hierarchical cluster tree (names as SIG1-SIG6 in Table 1) and allow us to capture the function of each cluster with a gene signature. For example, Chemokine-CLU is associated with a set of genes that function in G-protein signaling pathway and the response to endogenous and environmental insults, while TGF-CLU uniquely associates with another set of genes that mediates blood coagulations and plasminogen activating cascade which are often associated with the innate immunity in infectious and neuroinflammatory diseases (Table 1). We also identified two sets of genes (BLU1, BLU2) that are distant from the three major clusters (Chemokine-CLU, IFU-CLU, IL-CLU) (Table 1). These two distant genes have biological functions in the nucleus. The detailed gene list for each of these sets is listed in Supplementary Material Table S2.

**Figure 3.**
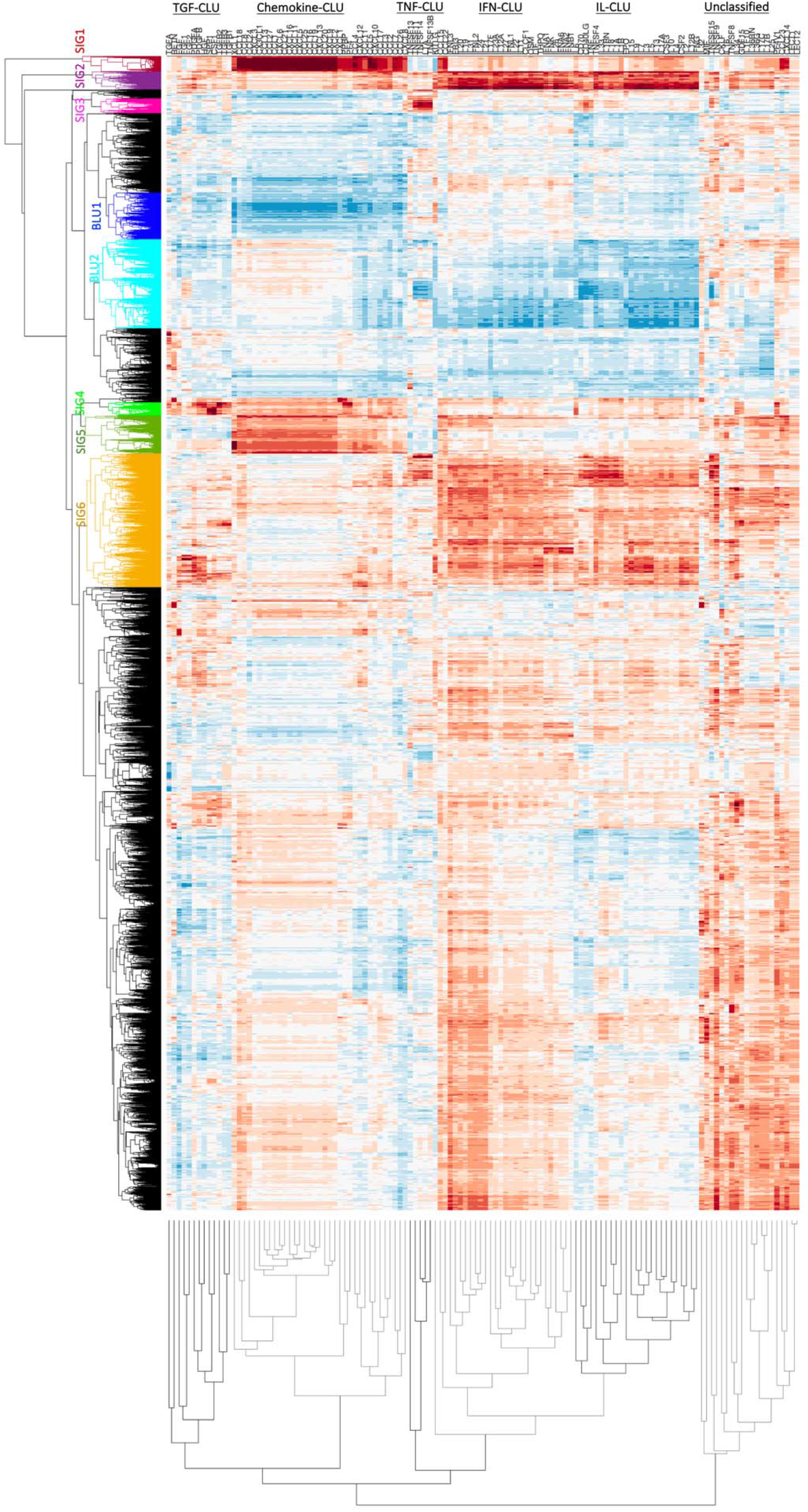
The 126 cytokines form six clusters based on their Association Scores with the 14,581 non-cytokine genes. The six clusters are named based on the most enriched types of cytokines: TGF-CLU (growth factors), Chemokine-CLU (chemokines), TNF-CLU (TNFs), IFN-CLU (interferons), IL-CLU (interleukins), and Unclassified-CLU. Based on the dendrogram of the hierarchical cluster tree, we identified six gene sets (SIG1-SIG6) that associate with the six individual clusters and two gene sets (BLU1, BLU2) that do not interact with the three major clusters (Chemokine-CLU, IFU-CLU, IL-CLU). Details of these signatures are in Table 1.

**Table 1.** Details of the six clusters in figure 3. The associations between the six gene sets (SIG1-SIG6) and specific clusters (CLU) are noted in (A). Specific cytokines in each of the six clusters are listed in (B). The functional annotations of the six genes sets (SIG1-SIG6) are in (C). The two gene sets (BLU1, BLU2) that do not interact with the three major clusters (Chemokine-CLU, IFU-CLU, IL-CLU) are listed in (D).

|  | TGF-CLU | Chemokine-CLU | TNF-CLU | IFN-CLU | IL-CLU | Unclassified-CLU |
| --- | --- | --- | --- | --- | --- | --- |
| SIG1 |  | Yes |  |  |  |  |
| SIG2 |  |  |  | Yes | Yes |  |
| SIG3 |  |  | Yes |  |  |  |
| SIG4 | Yes |  |  |  |  |  |
| SIG5 |  | Yes |  |  |  |  |
| SIG6 |  |  |  | Yes | Yes |  |

B. Cytokines in the six clusters
| TGF-CLU | Chemokine-CLU | TNF-CLU | IFN-CLU | IL-CLU | Unclassified-CLU |
| --- | --- | --- | --- | --- | --- |
| TGFA RETN EGF<br>FGF1 FGF2<br>PDGFA PDGFB<br>HGF SPP1 CSF1<br>TGFB2 TGFB3<br>TGFB1 | XCL1 CCL18 CCL8<br>CCL24 CXCL3<br>CX3CL1 CCL13<br>CCL27 CCL1<br>CXCL6 CCL21<br>CXCL16 CXCL11<br>CXCL5 CCL25<br>CCL16 CCL19<br>CXCL13 CCL20<br>CXCL9 CXCL2<br>CXCL1 PPBP PF4<br>CCL4 CXCL12<br>CCL11 CCL5<br>CXCL10 CCL17<br>CCL3 CCL2 CCL7<br>CCL26 CXCL8 | TNFSF13<br>TNFSF11<br>TNFSF14 LTA<br>TNFSF13B<br>KITLG | CCL15 CCL22<br>IFNL3 EBI3 IL20<br>IL19 IL31 IFNL2<br>IL24 IL21 IL17F<br>IL23A IL12A IL27<br>IFNL1 IL22 IL11<br>CLCF1 OSM LIF<br>THPO IFNA1<br>IFNK IL26 IFNA6<br>IFNA2 IFNB1 | IL6 CD70<br>CD40LG TNF<br>TNFSF4 IL33<br>IL1RN IL18 IL1A<br>IL1B EPO IL15<br>IL9 IL7 IL3 IL5<br>IL13 IL17A CSF3<br>IL10 IL4 CSF2 IL2<br>IL12B IFNG | IL32 MIF<br>TNFSF15 TNFSF9<br>CKLF IL16<br>TNFSF8 IL34<br>GDF15 IL1F10<br>IL36RN IL36G<br>IL17C IL17B IL25<br>PF4V1 CCL23<br>CXCL14 LECT1<br>LECT2 |

C. GO/PANTHER pathways of the six gene sets.
|  | GO/PANTHER Pathways |
| --- | --- |
| SIG1 | G-protein signaling pathways, inflammation mediated chemokine pathways, GABA-B receptor signaling, 5HT1 receptor signaling, Enkephalin release, Endothelin signaling, opioid pathways, dopamine pathways, Nicotine pharmacodynamics, Metabotropic glutamate pathways |
| SIG2 | Interleukin signaling, JAK/STAT signaling, Toll receptor signaling, Interferon-gamma signaling, inflammation mediated by cytokine signaling pathways |
| SIG3 | Ubiquitin proteasome pathways, Parkinson disease, Hedgehog signaling, WNT signaling, FGF signaling, Angiogenesis, hypoxia response via HIF activation, p53 pathways by glucose deprivation |
| SIG4 | Blood coagulation, Plasminogen activating cascade |
| SIG5 | G-protein signaling pathways, 5HT2 receptor signaling, Histamine H1 receptor signaling, Thyrotropin releasing hormone receptor signaling, Oxytocin receptor signaling, Muscarinic acetylcholine receptor signaling, Corticotrophin releasing factor receptor signaling, Angiotensin II stimulate signaling, Endogenous cannabinoid signaling |
| SIG6 | Apoptosis signaling, Toll receptor signaling, FAS signaling, p38 MAPK pathway, Gonadotropin-releasing hormone receptor pathways, B-cell activation, T-cell activation, Insulin/IGF pathway (protein kinase B cascade, MAPKK/MAPK cascade), axon guidance mediated by netrin, Angiogenesis, EGF signaling, PDGF signaling, VEGF signaling |

D. GO/PANTHER pathways of gene sets that are distant from immune responses.
|  | GO/PANTHER Pathways |
| --- | --- |
| BLU1 | Ribosome biogenesis, rRNA metabolic process, gene expression, translation |
| BLU2 | DNA repair, DNA metabolic process, telomere maintenance, cellular macromolecule metabolic process, response to stress |

### The predicted cytokine profiles of 171 diseases are validated using literature

We have collected gene sets associated with 11,944 diseases from DisGeNET ^21^. A majority (5048) of these diseases are linked with fewer than ten genes, while a few are associated with up to 2000 genes (Supplementary Material Figure S4-S6).

For each disease, we estimated the normalized average Association Scores (NAAS) between each cytokine and the disease based on the normalized score of averaged Association Scores of the given gene set (Figure 1B), resulting in a 126-dimension (the embedding dimension) cytokine profile for each disease. From the 11,944 diseases, we identified 171 well-studied diseases of which cytokine associations (for 79 of our 126 essential cytokines) have been evaluated by their co-occurrence frequency in literature sampling in ImmuneXpresso. The predicted profiles correlate with the literature sampling frequency significantly (p-value <0.05) for 147 of the 171 diseases, suggesting reasonable reliability of predicted cytokine profiles (Figure 4A, Supplementary Material Table S3). Figure 4B shows an example of the predicted cytokine profiles for aneurysm, a disease that is not typically considered as an immune disorder. The NAAS between aneurysm and each of the 79 cytokines of which the literature sampling frequency in diseases are known in ImmuneXpresso are plotted with known associations (frequency cutoff is 0.005) marked in solid blue squares. At a high cutoff (NAAS >0.8), the recall rate is 21/24. The novel predictions are: HGF, IL11, IL12B, IL13, IL15, IL17F, IL22, IL33IL5, IL7, IL9, LIF, OSM, PDGFA, PDGFB, PPBP.

**Figure 4.**
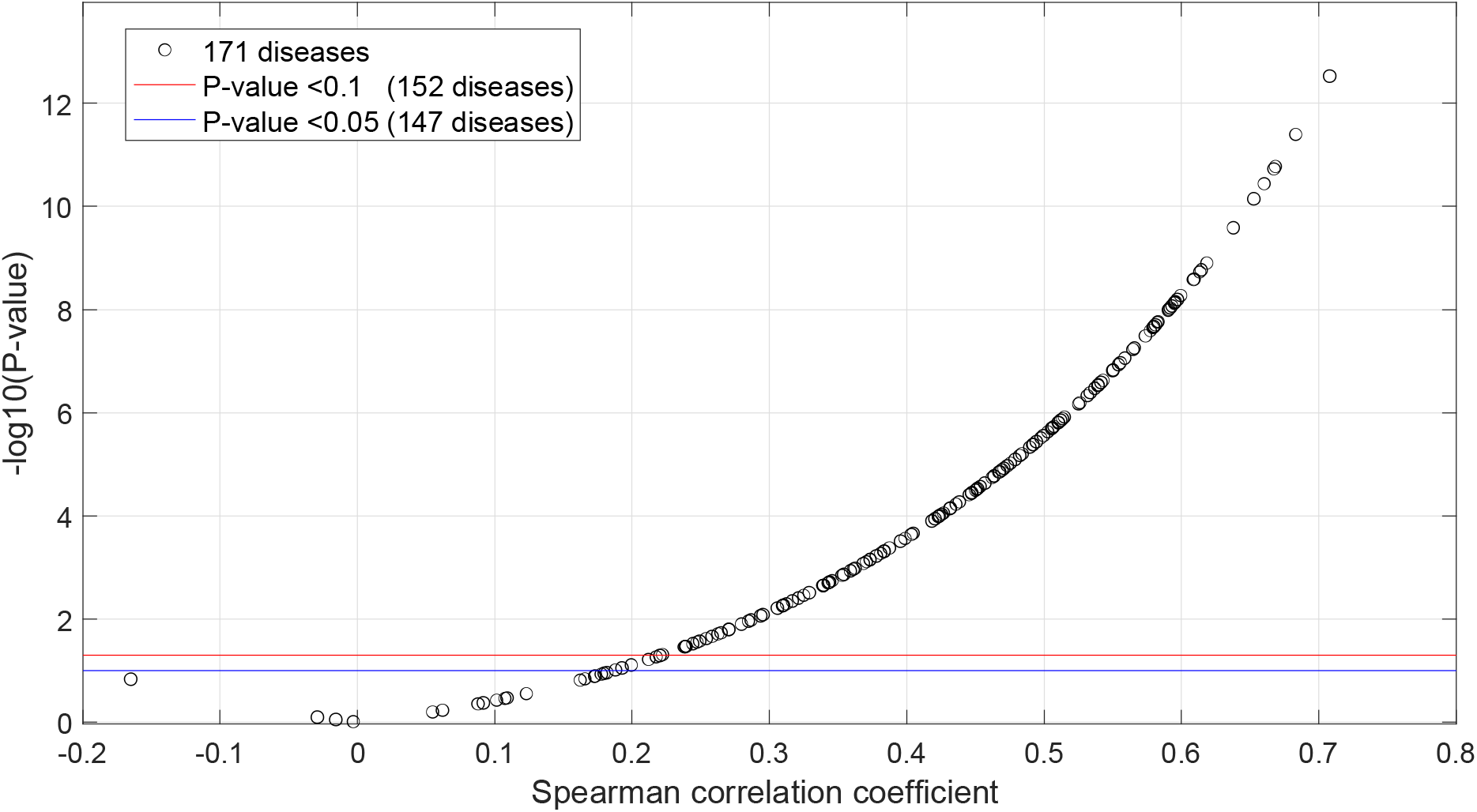
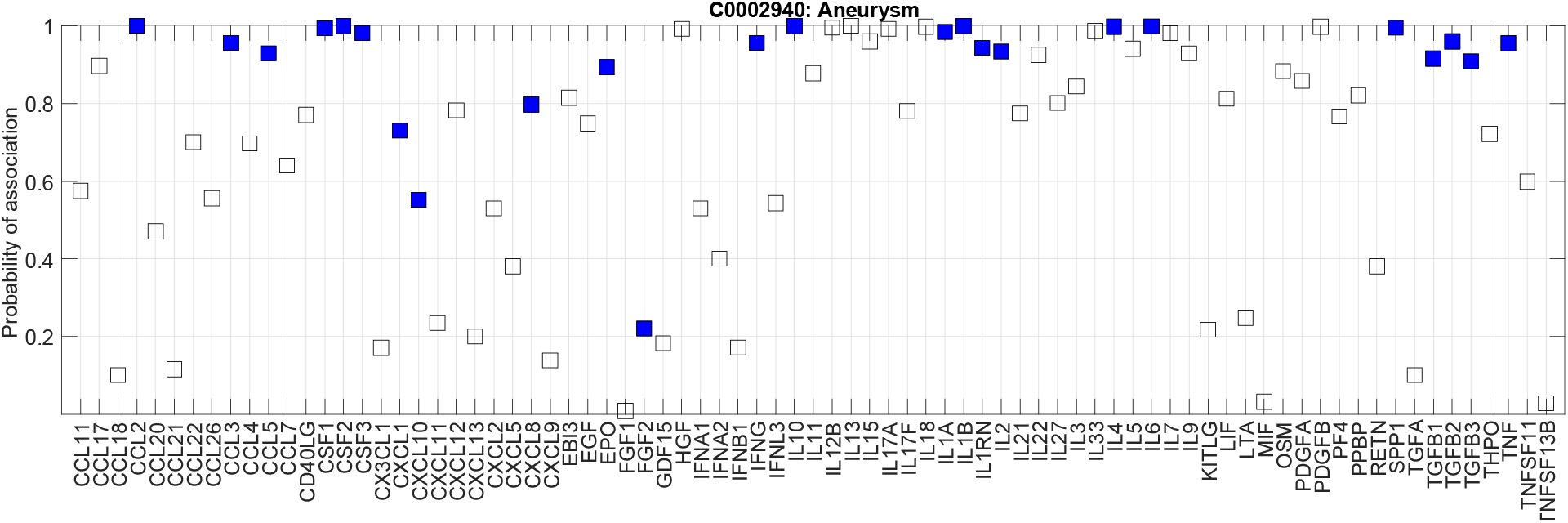
A. Predicted cytokine profiles for 171 well-studied diseases correlate with cytokine sampling in literature. Given one disease, we estimated the Probability of Association between each cytokine and the disease based on the normalized score of averaged Association Scores of the given gene set, resulting in a 126-dimension cytokine profile for each disease. The Spearman correlation coefficients between the predicted Probability of Association and the know literature sampling frequencies are plotted against the P-values. Of the 171 diseases, we were able to predict 147 diseases with p-value<0.05, suggesting the accuracy of the predicted profiles. B. The Probability of Association between aneurysm and each of the 79 cytokines of which the literature sampling frequency in diseases are known in ImmuneXpresso. Known associations (frequency cutoff of 0.005 in ImmuneXpresso) are marked in solid blue squares.

### Defining the key modes of cytokine response

We have shown that the predicted disease profiles of 79 cytokines align with known cytokine associations, suggesting the reliability of disease profiles of 126 cytokines. There are many ways to classify diseases, we aimed to assess patterns in cytokine response for different diseases, reasoning that shared cytokine response might indicate the potential for shared treatment strategies. We analyzed the predicted disease profiles of 126 cytokines in the 171 well-studied diseases. The 171 diseases include 23 immune disorders (C20), 48 infections(C01), seventeen cardiovascular diseases (C14), thirteen metabolic disorders(C18), and 55 neoplasms (C04). Note that one disease may belong to more than one disease classes (Supplementary Material Table S4). They fall into three distinct patterns in cytokine responses (Figure 5A). The majority of metabolic diseases (11/13) and cardiovascular disorders (11/17) are enriched in one cluster, named cluster-1 (blue in Figure 5A), suggesting that cytokine responses are shared across different disease classes. Immune disorders are enriched in cluster-3. Infections are split into two clusters (cluster-2, green and cluster-3 orange in Figure 5A). Note that of the twenty neoplasms in cluster-3, nineteen are hematic and lymphatic diseases (C15/C04), suggesting that these neoplasms have distinct cytokine distributions from other neoplasms. The three themes of cytokine responses across five diseases class suggest the common framework shared by different disease classes, and the heterogeneity of cytokine responses within a disease class.

**Figure 5.**
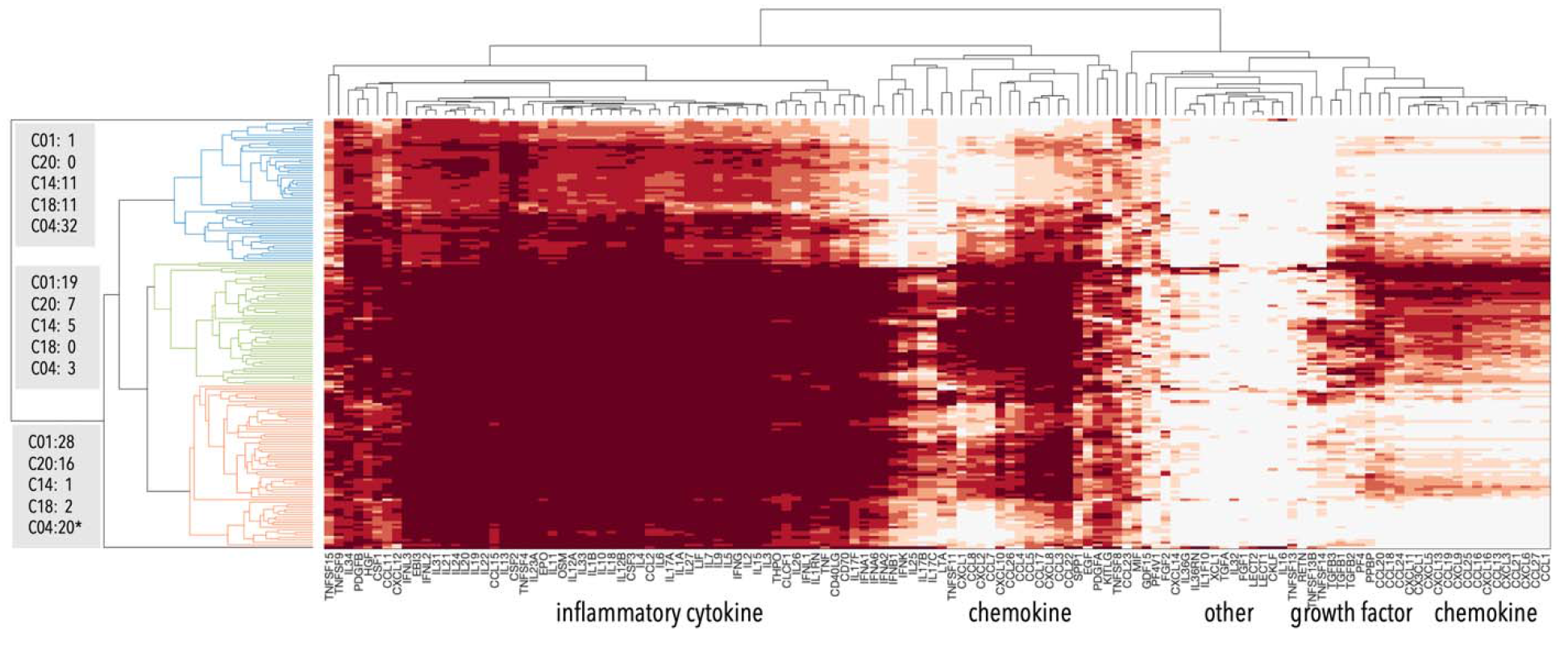
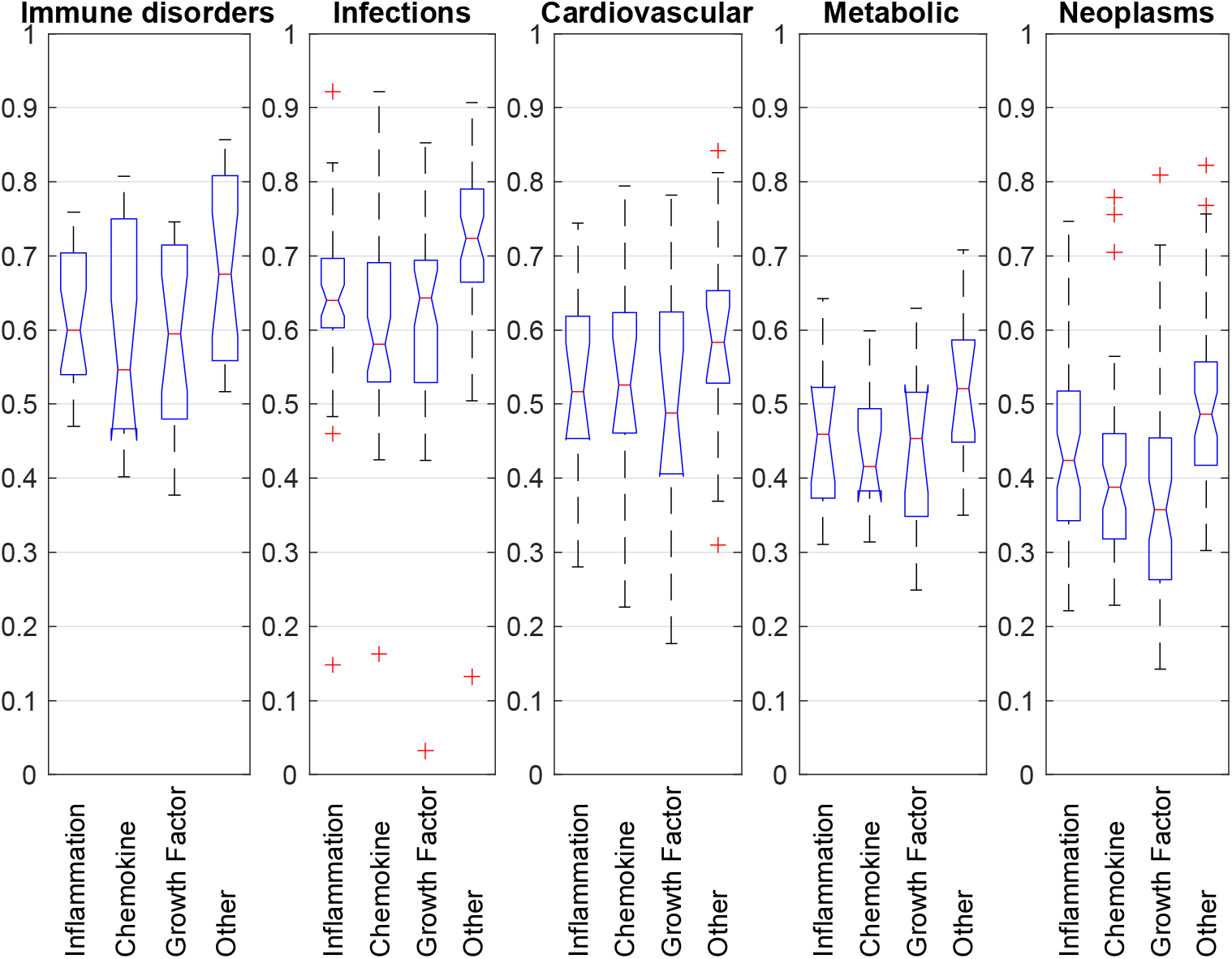
A. Cytokine features for the 171 well-studied diseases. The 171 diseases formed three clusters based on their NASS with different types of cytokines. Infections (01) dominant in cluster-1 (red and orange). Note that of the twenty neoplasms in cluster-1, nineteen are hemic and lymphatic diseases (C15/C04). Cluster-2 (green) represents a mix of immune disorders (C20) and infections (C01). Cluster-3 (blue and cyan) represent a mix of cardiovascular disease (C14), metabolic diseases (C18), and neoplasms (C04). Note that diseases of other classes are not counted in the labels. Cluster details are in Supplementary Material Table S4. B. Immune Scores of five disease classes (23 immune disorders, 48 infections, 17 cardiovascular, 13 metabolic, 55 neoplasms). For each disease within a class, the average of NAAS between disease and the cytokines in four categories are plotted: 47 inflammation related cytokines, 37 chemokines, 13 growth factors, and 29 other cytokines are plotted. The chemokine scores for immune disorders spread in a wide range. While for infections, growth factors have the highest scores, compared to immune disorders and the other three disease classes. Cardiovascular diseases have higher scores than metabolic diseases over the three groups of cytokines. Neoplasms show the lowest scores for all four categories.

We observed patterns of interacting with genome in different families of cytokines (Figure 3). The disease cytokine profiles show that inflammatory cytokines that include interleukins, inferons, TNFs are grouped together (Figure 5A, horizontal), while chemokines form two groups based on their associations with diseases. These groups of cytokines decide the clustering of diseases across different classes. Therefore, we analyzed the influence on diseases from inflammatory components, chemokines, growth factors, and other cytokines. We define Immune Scores to capture the contributions from four categories of cytokines to the inflammation process of a given pathogenesis (Figure 5B). The chemokine scores of immune disorders spread in a wide range. Infections have the highest scores for growth factors. Cardiovascular diseases have higher scores than metabolic diseases in all four categories, while neoplasms show the lowest scores in all the categories. In summary, a disease can be represented by a 126-dimension cytokine profile or a 4-dimensional summary Immune Scores, both of which suggest that the inflammatory responses in different diseases are mediated by different distributions of cytokines.

### Inflammatory response subnetworks provide disease-specific insight

To gain a more comprehensive understanding of inflammatory components, we identified the subnetworks formed by the predicted cytokines and pathogenesis genes of a given disease. We inspected the subnetworks of five diseases representing immune disorders, infections, cardiovascular diseases, metabolic disorders, and neoplasms (systemic lupus erythematosus (SLE), TB, aneurysm, metabolic syndrome X, and acute leukemia) and found that they visually show different network patterns (Supplementary Material Figure S8). We quantified the network features by counting the number of high confidence associations between disease-associated genes (DisGeNET) and cytokines in the five example diseases (Table 2A). For these five diseases, 8-14% of the known disease-associated genes in DisGeNET are predicted to interact with at least one of the 126 cytokines, suggesting that the connections between pathogenesis and immune response are less dense compared with metabolic or signaling modules.

**Table 2.**
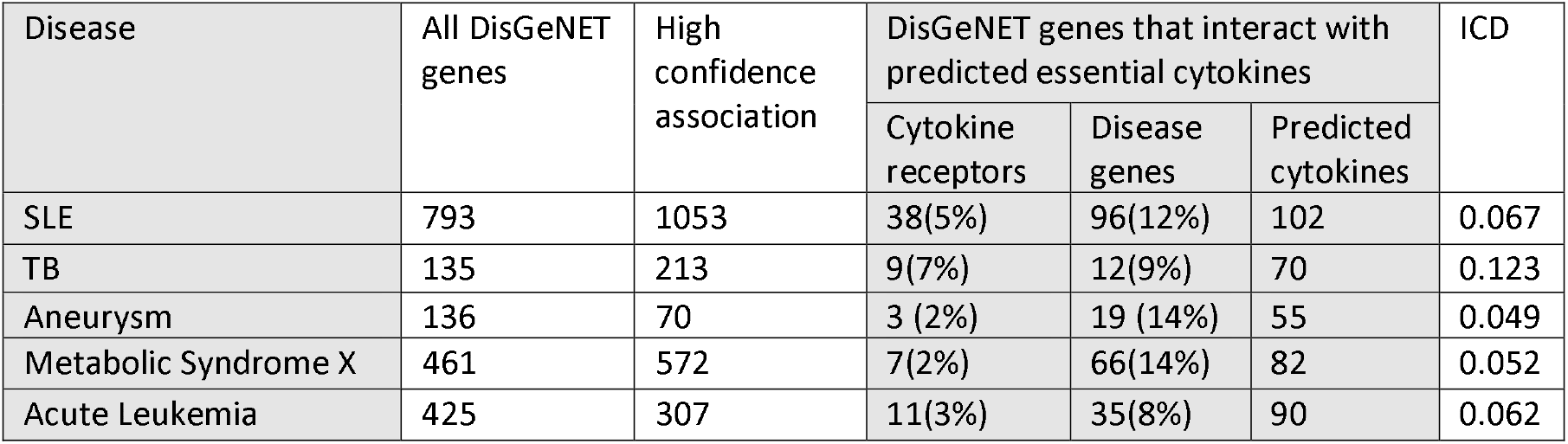
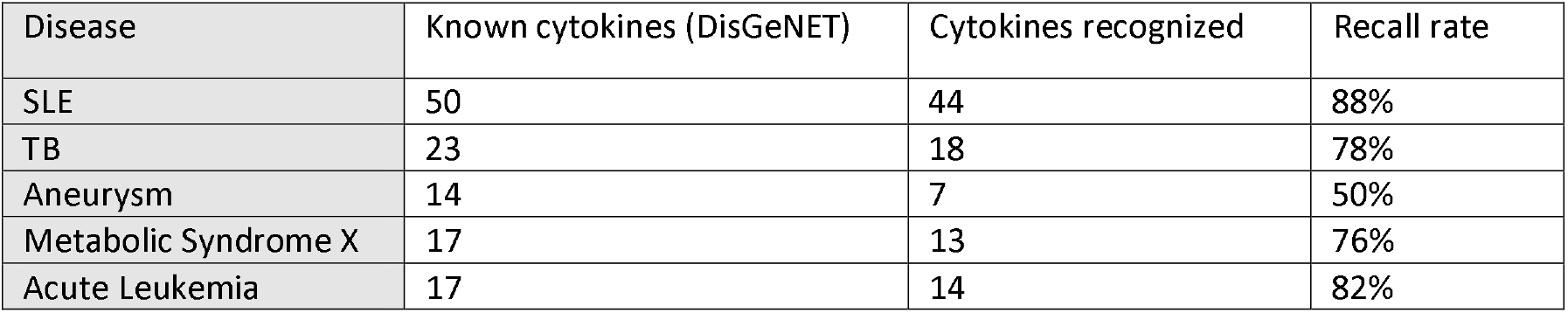
Analysis of subnetworks formed by high confidence associations (Association Score cutoff as 0.8) between the known disease associated genes (DisGeNET) and the predicted cytokines in five diseases. A. The known DisGeNET genes (column #2) of a given disease often contain cytokine receptors and other genes. The numbers of cytokine receptors and other disease genes captured by the high confidence associations (column #3) are in column #4 and column #5, respectively. The numbers of the predicted essential cytokines that interact with receptors and disease genes from DisGeNET are listed in column #6. The Immune Connection Density (ICD) estimated on the subnetworks formed by receptors, disease genes and essential cytokines for each disease are in column #7. B. The known DisGeNET genes of a given disease often contain cytokines already. The recall rate (column #3) of these cytokines being recognized by the predicted subnetworks formed by high confidence associations ranges from 50% to 88%.

| Disease | All DisGeNET genes | High confidence association | DisGeNET genes that interact with predicted essential cytokines |  |  | ICD |
| --- | --- | --- | --- | --- | --- | --- |
|  |  |  | Cytokine receptors | Disease genes | Predicted cytokines |  |
| SLE | 793 | 1053 | 38(5%) | 96(12%) | 102 | 0.067 |
| TB | 135 | 213 | 9(7%) | 12(9%) | 70 | 0.123 |
| Aneurysm | 136 | 70 | 3 (2%) | 19 (14%) | 55 | 0.049 |
| Metabolic Syndrome X | 461 | 572 | 7(2%) | 66(14%) | 82 | 0.052 |
| Acute Leukemia | 425 | 307 | 11(3%) | 35(8%) | 90 | 0.062 |

| Disease | Known cytokines (DisGeNET) | Cytokines recognized | Recall rate |
| --- | --- | --- | --- |
| SLE | 50 | 44 | 88% |
| TB | 23 | 18 | 78% |
| Aneurysm | 14 | 7 | 50% |
| Metabolic Syndrome X | 17 | 13 | 76% |
| Acute Leukemia | 17 | 14 | 82% |

The genes involved in the pathogenesis of a given disease often are cytokine receptors – and these are critical clues for how the pathogenesis genes may communicate with immune modules. The numbers of cytokine receptors that interact with cytokines suggest the different density of inflammatory responses. For example, cytokine receptors are more heavily involved in the process from pathogenesis to inflammatory responses in SLE and TB, but not in aneurysm, metabolic syndrome X and acute leukemia. In order to capture the density of connections between pathogenesis genes and immune response, we compute a “Immune Connection Density (ICD)”. We adapted the original equation of network efficiency ^22^ by counting the edges between the two components within the predicted subnetworks: genes for pathogenesis and for inflammation (see Methods). The interactions between cytokines, or between pathogenesis genes are not included in the calculation. Therefore, the ICD suggests the associations between the two functions, pathogenesis and inflammation processes. Compared to SLE, aneurysm, metabolic syndrome X, and acute leukemia, TB shows the highest ICD (Table 2A), suggesting coherent reactions of immune systems upon infections.

Analyzing the subnetworks formed by the predicted cytokines and pathogenesis genes also suggests the reliability of the predicted disease profiles of 126 cytokines (See Result section “Cytokine features of diseases”). Some of the known genes from DisGeNET are cytokines. When comparing with all the cytokines known in DisGeNET, the recall rate (Table 2B) of being recognized by the high confidence interactions ranges from 50% to 88%. When using lower confidence cutoff (0.7), 36 more essential cytokines are predicted to be associated with aneurysm, and 11/14 (78%) of the known essential cytokines can be recognized by our network methods.

### Immune disorders pathogenesis genes connect to key inflammatory genes

Within the class of immune disorders, we observed different detailed predicted cytokine associations in individual diseases (Figure 6). The five diseases, rheumatoid arthritis (RA), psoriasis (PS), ulcerative colitis (UC), Crohn’s disease (CD) and SLE show similar associations with the core cytokines for inflammation, but differ in their detailed associations with chemokines, TNFs and growth factors.

**Figure 6.**
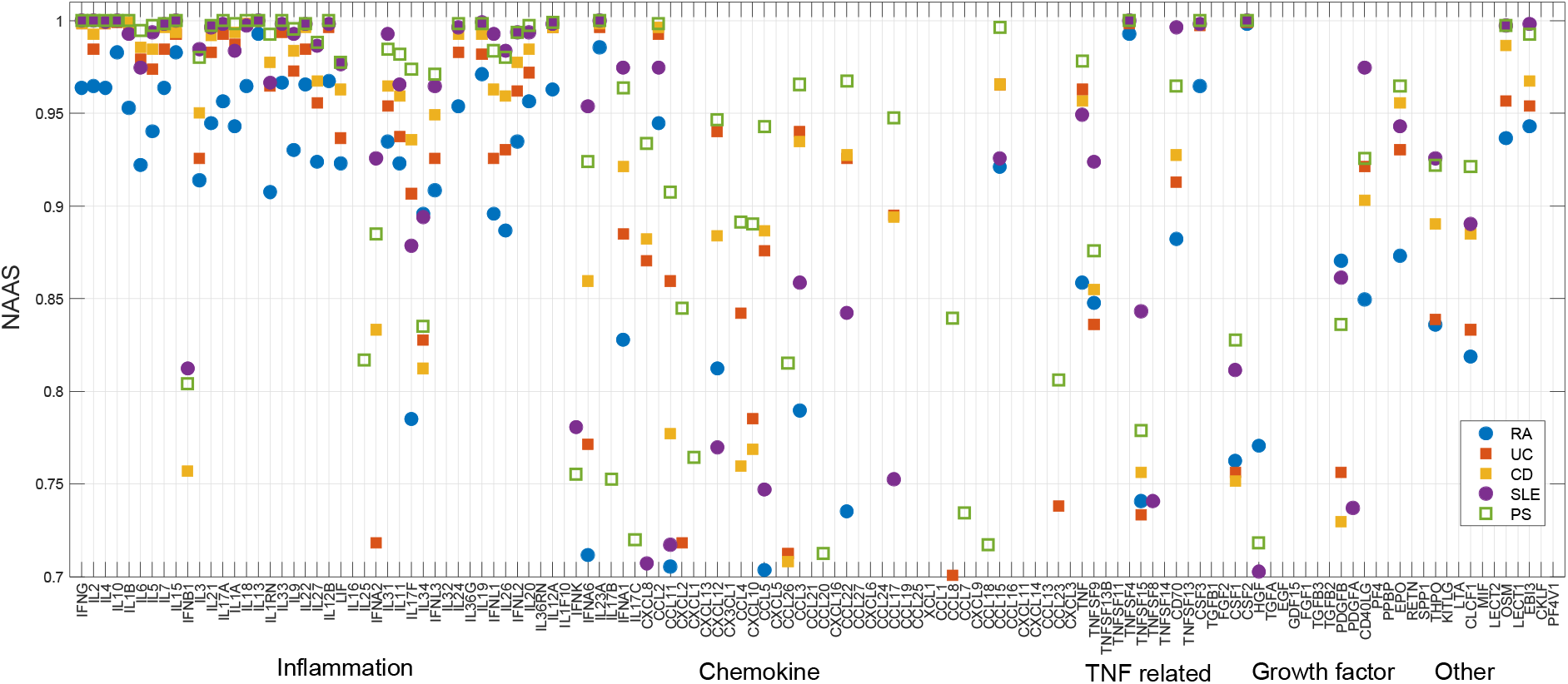
Disease-specific cytokine profiles of five immune disorders. Y-axis shows the Probability of Associations between cytokines and five immune disorders: rheumatoid arthritis (RA), psoriasis (PS), ulcerative colitis (UC), Crohn’s disease (CD) and systemic lupus erythematosus (SLE). Conserved association pattern is observed in inflammation related cytokines, while differential patterns are observed in other types of cytokines.

In order to understand how pathogenesis genes associate with different types of cytokines, we used a force-directed graph to visualize the various interactions in the process from pathogenesis to inflammatory responses under different disease contexts. Stronger associations are drawn with stronger “springs” in order to convey a qualitative understanding of the subnetwork structure ^23^. The visualization in Figure 7 shows that the individual pathogenesis genes of SLE link to their embedded network neighbors--shorter edges correspond to closer network neighbors and thus allows us to assess the pathogenesis genes that are most likely associated with inflammation. Different sets (Box-C and Box-I-1-4) of pathogenesis genes form associations with chemokines and pre-inflammatory cytokines, respectively. Six genes (ACKR3, HRH4, HTR1, GAL, GRM3, S1PR1) are connected with a set of densely connected chemokines (23 cytokines marked in red box), with ANXA1 interacting with sixteen chemokines. Seven pathogenesis genes (Box-I-1) make interactions with a group of 28 pre-inflammation cytokines (orange box) directly or through receptors (green nodes). Next to this group of pre-inflammation cytokines, a group of nine disease associated genes (Box-I-3, S100A8, NLRP3, MYD88, IRAK1, IRAK4, TIRAP, TLR2, TLR5, TLR9) form a small clique with four other pre-inflammation cytokines (IL18, IL22, IL1A and IL1B). Two other groups of genes (Box-I-2: PSME3, PSMB9, PSMD4, PSMB6, PSMA5, PSMD7, OAZ1, REL, HIVEP3, and Box-I-4: MAP4K3, SPATA2, TNIP1, ZC3H12A) are distant from these pre-inflammation cytokines (orange dots), even they are apparently linked to the TNFs.

**Figure 7.**
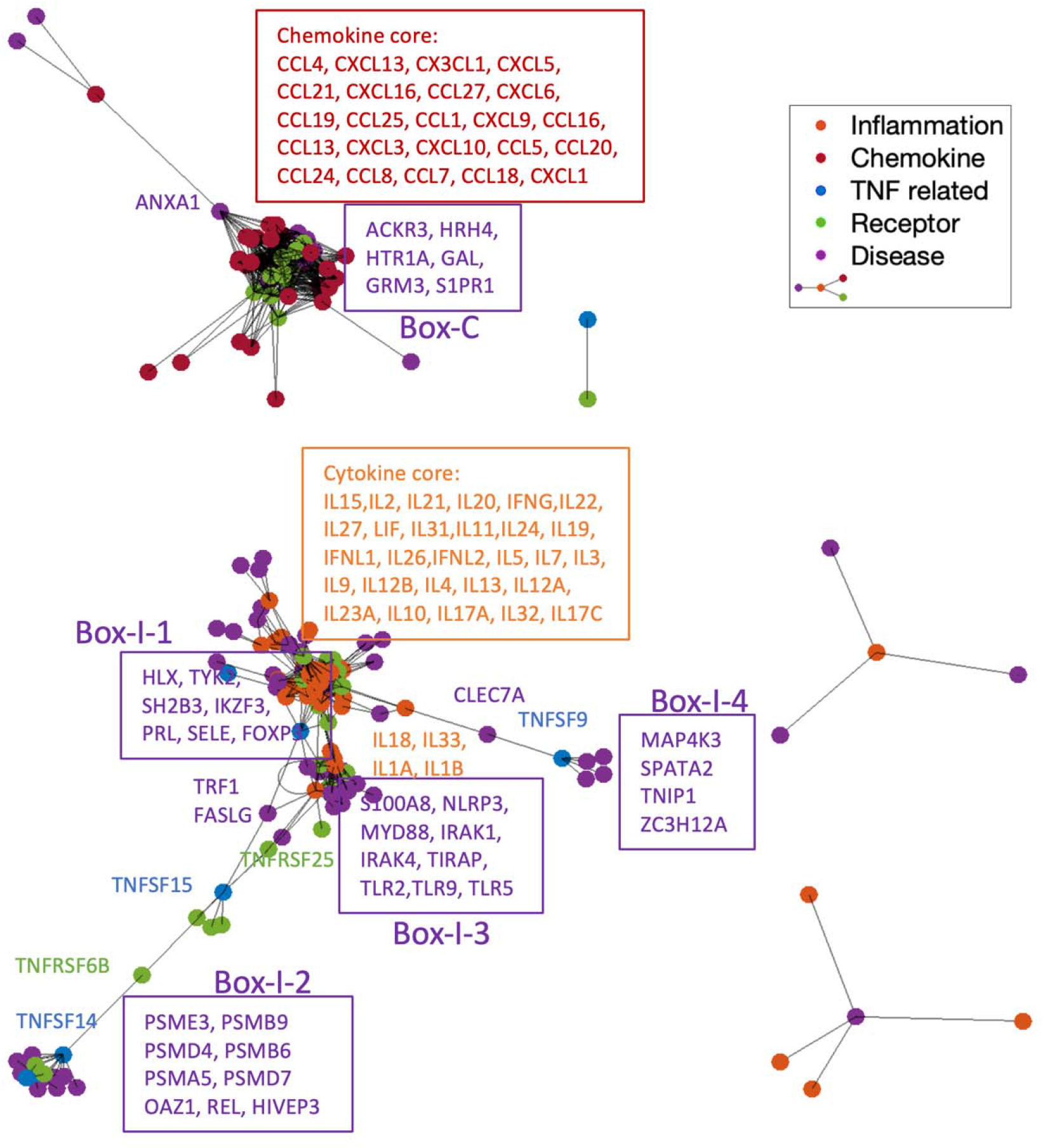
SLE subnetwork formed between pathogenesis genes (green and purple) and inflammatory responses (orange, red, blue). The graph was plotted in force-directed layout that uses attractive forces between adjacent nodes and repulsive forces between distant nodes. The distances between two vertices are roughly proportional to the length of the shortest path between them. Six genes (ACKR3, HRH4, HTR1, GAL, GRM3, S1PR1) in Box-C are making high degree contacts with chemokine core (red box), with ANXA1 interacts with sixteen chemokines. Interactions with inflammation core (orange box) form multiple directions. Seven pathogenesis genes (Box-I-1) are making interactions with core of inflammation cytokines (orange box) directly or through receptors (green nodes). Nine disease genes in Box-I-3 form a small core with four cytokines (IL18, IL22, IL1A and IL1B). Two other groups of genes (Box-I-2 and Box-I-4) stay remotely from cytokine cores, even they are apparently linked to the TNFs, as they cannot overcome the repulse force to association with the center of inflammation responses.

### Highly connected graphs capture connections between immune disorder pathogenesis and inflammation

The ICD for the five immune disorders are: RA, 0.055, UC, 0.069, CD, 0.71, SLE, 0.067, and PS, 0.060, suggesting that density of association between pathogenesis and inflammation are similar. However, different sets of genes are involved in these interactions. For each specific immune disorder, we are interested in the key pathogenesis genes that mediate cytokine connections. From the predicted inflammatory response subnetworks, spectrum partition enabled identification of highly connected graphs, or modules formed by a group of well-connected cytokines and pathogenesis genes, revealing the key genes for cytokine mediators that drive the pathogenesis in inflammatory diseases. Table 3 shows that 23 receptors and 36 disease genes (from 1340 known genes) were identified to form well-connected cytokine modules in RA); six receptors and four disease genes (from 542 known genes) for PS, seventeen receptors and 35 disease genes (from 793 known genes) for SLE, four receptors and eight disease genes (from 654 known genes) for UC, and thirteen receptors and eighteen disease genes (from 622 known genes) for CD. This process further prioritizes the known disease associated genes from DisGeNET, providing a more focused set of candidates for experimental follow up. In addition, the cytokine modules identified for SLE and RA overlap, while those identified in PS and UC overlap, but not the corresponding pathogenesis genes (Table 3 and Supplementary Material Table S5), suggesting different mechanisms mediating the cytokine framework.

**Table 3.**
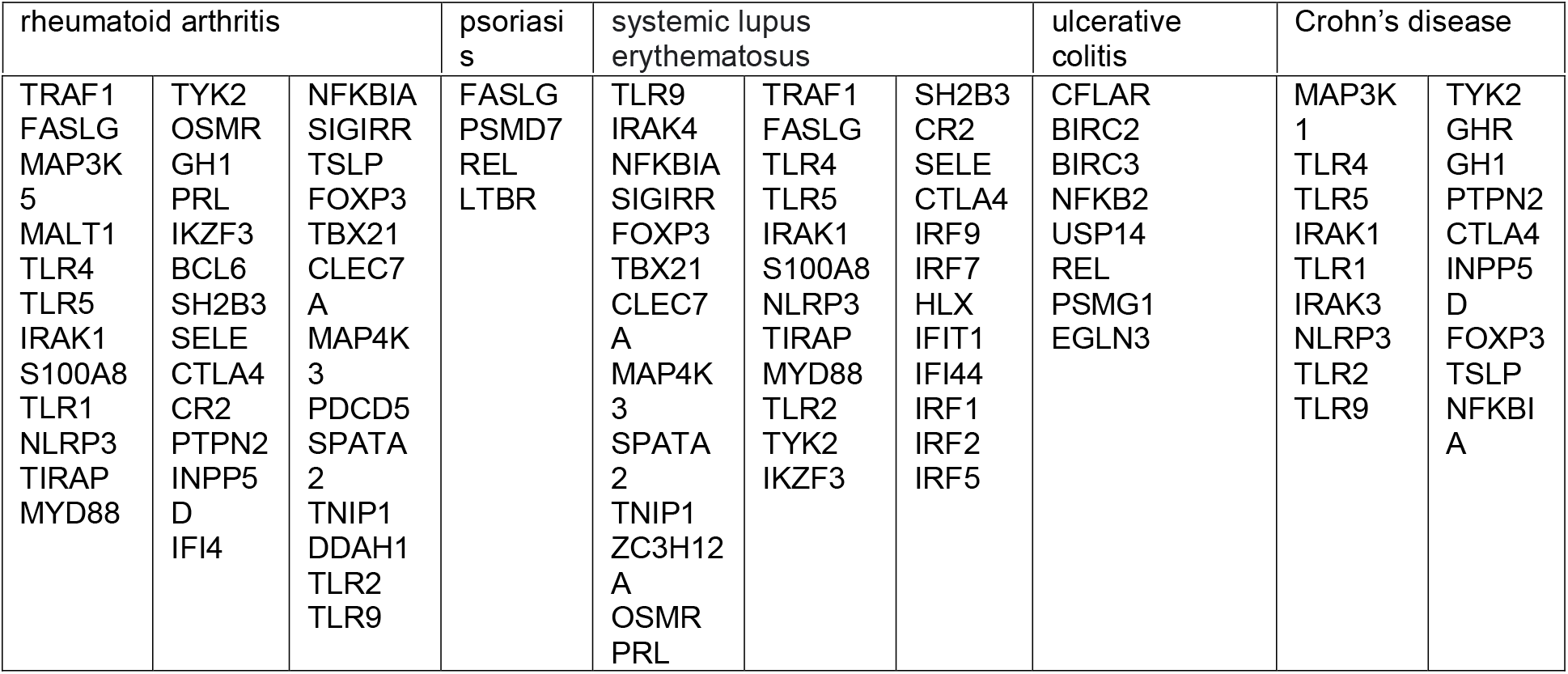
Pathogenesis genes in the highly connected modules identified by spectrum partition on the subnetworks formed by pathogenesis genes, receptors, and cytokines, in the context of five immune disorders: rheumatoid arthritis (RA), psoriasis (PS), ulcerative colitis (UC), Crohn’s disease (CD) and systemic lupus erythematosus (SLE). From the 1340 disease associated genes for RA, 36 disease genes are identified in the cytokine modules (2.7%). Correspondingly, four disease genes from the 542 genes for PS (0.7%), 35 disease genes from the 793 genes for SLE (4.4%), eight disease genes from the 654 genes for UC (1.2%), and 18 disease genes from the 622 genes for CD (3%).

## Discussion

### A complete map of associations between genes in large scale enables the identification of genome-wide features of cytokine interactions

The connection rates between immune response genes are lower than that within classic functional modules heavily studied modules for metabolic, transcriptome, signaling. Meanwhile, the associations between functional units are not well defined, compared with the connections within known functional units (Supplementary Materials Table S1). Disease associated genes are often found scattered in different modules (metabolic, signaling and immune modules). For example, for non-alcoholic steatohepatitis, disease-associated genes are found in the highly disparate functional modules of fatty acid beta-oxidation, proteolysis, signal transduction, leukocyte aggregation and other cellular process^14^. In this work, we aimed to identify novel associations between pathogenesis genes and immune responses; for this task, we required a map of pairwise associations between genes on a large scale. The STRING network itself is a highly connected graph that obeys “small world” statistics and thus path length calculations are not useful for estimating likelihood of association^18^. The embedding space provides more information by capturing the topological structures of STRING, enabling a complete map of pairwise associations.

The wide pleiotropy and the element of redundancy in cytokine family (each cytokine has multiple functions, and each function potentially mediated by more than one cytokine) make the classification of cytokines a challenge^20 24^. Using our map of pairwise associations, we were able to connect a key set of cytokines to 14K human genes and identified genome-wide features that interact with unique groups of cytokines. This genome-wide associations enable a more systematic classification of cytokines. The biological annotations of these specific interactions also provide important insights into the functions of cytokine groups. We identified two sets of genes (191 genes in SIG1 and 175 genes in SIG5) that interact only with chemokines, highlighting specific signaling pathways for chemokines: their biological functions focus on G-protein signaling pathways, the response to endogenous and environmental insults. We also found that genes responsible for Ubiquitin proteasome pathways interact with TNFs, not other cytokines (SIG3 in Table 1). Another interesting set of gene is SIG4 that interacts only with TGF and plays important roles for blood coagulation and plasminogen activating cascade, which are often associated with the innate immunity in infectious and neuroinflammatory diseases. Some of the genes in the featured interactions (F5 and SERPINE2 in SIG4) are known to affect the concentrations of circulating cytokines ^25^. Those genes that are not recognized by GWAS could be critical links from pathogenesis to inflammation. The biochemical pathways underlying the links from these genes to complex diseases have remained elusive. Our findings provide candidate genes pivoting to deeper studies of pathogenesis and inflammation.

### Disease-specific cytokine profiles reveal flexible features of differential inflammatory responses

Our analysis suggests that diseases have flexible cytokine distributions even though there may be a shared cytokine framework that provides conserved mechanisms of inflammation.

First, clustering diseases based on their cytokine profiles shows three different cytokine response modes, these clusters correlate with disease classification. For example, of the 55 neoplasms, 32 fall into cluster-1 and twenty in cluster-3, of which nineteen are hemic and lymphatic diseases (C15/C04) (Figure 5A). Second, the Immune Scores that capture the contributions from different types of cytokines to the inflammation show that inflammation as a driver of pathology for many diseases beyond those that are typically considered autoimmune and infectious. Cardiovascular diseases show higher scores than metabolic disorders and neoplasms (Figure 5B). The increased concentrations of cytokines in cardiovascular disease are not only markers of chronic low-grade inflammation, but also provide an important pathophysiological link between cardiovascular health and ageing^26^. Third, the numbers of disease genes and receptors that are associated with essential cytokines vary widely compared with the numbers of cytokines themselves, suggesting different mechanisms mediate between pathogenesis and inflammatory response (Table 2). Finally, the ICD quantifies the density of interactions between pathogenesis and inflammation processes and suggests the mechanism by which different diseases have different levels of inflammation (Table 2).

Within the class of immune disorders, we also observed differential cytokine distributions between different diseases (Figure 6). The five immune disorders show close interactions with the cytokines responsible for pre-inflammatory responses, but not all five of them have close interactions with chemokines, TNFs or growth factors. One explanation is that multiple cytokines are triggered simultaneously by a few key activated triggers. Therefore, identification of the key genes and cytokines that trigger the immune responses in individual diseases may provide insights into therapeutic strategies.

### Subnetworks between pathogenesis and inflammation suggest different mechanisms of immune response

Our predicted associations between cytokines and pathogenesis enable network analysis from different perspectives, providing useful insights into the molecular pathways that mediate inflammation. We investigated two methods for visualizing the subnetworks formed between pathogenesis and inflammation. Through the hierarchical layered analysis on the connections between pathogenesis and inflammation, we were able to identify the central nodes in this dynamic process ^27^ (Supplementary Materials Figure S9). For example, in the subnetwork of in metabolic syndrome X, pathogenesis genes AGTR2 and ADRA1A are at the top hierarchical layer for chemokine signaling, and IRF1, MXZB1, MTTP and CNTC are at the top layer in cytokine signaling. In addition, the layered graph analysis suggests different interaction patterning: aneurysm showed a clear hierarchical flow starting from disease genes to cytokines, while metabolic syndrome X showed interactive layers between disease genes and cytokines, with an emphasis on chemokine responses, suggesting different mechanisms in signaling between pathogenesis and inflammation. We also adapted a force-directed graph to present the various interactions under different disease contexts ^23^ (Supplementary Materials Figure S8). The networks in SLE and TB display different patterns, suggesting different mechanisms in triggering inflammatory responses in immune disorders and infections, which may be related with the speed or strength of the immune reactions. Attractive forces between pathogenesis and chemokine responses are prominent in metabolic syndrome X, but not in aneurysm or acute leukemia. Interestingly, recent research found that modification in the genes that closely interact with chemokines may affect functions in glucose and lipid metabolism in patients with metabolic syndrome X ^28 29^. Our subnetwork of metabolic syndrome X provides candidates as novel targets of broader and more efficacious treatments and prevention of metabolic disease.

### Spectrum partition of subnetworks identifies key mediators of immune disorders

Immune cells can release many pathogenic cytokines. Mechanistic studies will be necessary to identify the key cytokines for a given inflammatory disorder and to pinpoint which cytokines might be the appropriate targets for tacking each disease. Given a disease, our methods identify the well-connected subnetwork formed between pathogenesis and inflammation and extract key genes closely associated with cytokines within the subnetwork. These key genes serve as candidates of therapeutic targets, as they are the main mediators to fuel inflammation. The human trials targeting different cytokines suggest the existence of a hierarchical framework of cytokines that defines groups for chronic inflammatory diseases rather differently from the homogenous molecular disease pattern previously assumed^12^. Interestingly, we have observed a common well-connected subnetwork that defines the close interactions between pathogenesis genes and cytokines in SLE and RA, which comprises pathogenesis genes TNIP1, SPATA2, MAP4K3, and CLEC7A. Annotation of these genes explains the possible shared pathways in SLE and RA, therefore shared therapeutic targets. Among these genes, TNIP1 is involved in inhibition of nuclear factor-κB (NF-κB) activation by interacting with TNF alpha-induced protein 3, an established susceptibility gene to SLE and RA ^30^. Other evidence suggests that the downregulation of SPATA2 augments transcriptional activation of NF□κB and inhibits TNF□α□induced necroptosis, pointing to an important function of SPATA2 in modulating the outcomes of TNF□α signaling, which plays important roles in inflammatory responses in RA and SLE ^31^. These observations validate our predicted key mediators for pathogenesis and inflammation. Further study on other key genes identified from disease-specific subnetworks may provide more insights into therapeutic strategies.

## Methods

### Network embedding of 14K human genes

We downloaded the network of 19,344 human genes from the STRING database which contains 5,879,727 edges. We filtered 14,707 genes that involve 728,090 high confidence edges (at a cutoff of 800) in STRING (Supplementary Material Table S1). We apply the methods of network scalable feature learning ^19^ that captures network topology features of the 14,707 genes in a 64-dimensional embedding space. Specifically, we have conducted an experiment to grid search the optimal hyperparameter for the embedding algorithm. The finalized parameters are: for each node, we use it as source to sample ten paths, with each path at a length of thirty (We set the hyperparameters as length of walks=30, number of walks=10, min count=1, batch word=6, window=10.). We then fit this data into node2vec algorithm to get the 64-dimensional embedding representation for each node.

### Prediction of Association Scores

Given any two genes from the pool of 14,707 human genes, we calculate the cosine distances between two embedded vectors, resulting in scores of possible pairs (108,140,571 pairs) between the 14,707 human genes, referred to as Association Scores. Among these pairs, the confidence scores of 9,250,034 pairs are available in STRING, of which 8,521,944 pairs have confidence scores below 800 and were not used for embedding. We evaluated the prediction of Association Scores by comparing with the known confidence scores for these 8,521,944 pairs.

### Prediction of disease-specific cytokine profiles

We identified 126 cytokines by mapping cytokines from ImmuneXpresso with the 14,707 human genes. For each cytokine gene, we calculated its Association Scores with the other 14,581 non-cytokine human genes. We have collected gene sets associated with 11,944 disease concepts from DisGeNET ^21^. For each disease concept, we calculated the average Association Scores between genes and each of the 126 cytokines, resulting in a 126-dimension cytokine profile for the disease. The 11,944 diseases were grouped into four bins based on the number of genes associated with the disease: 2-9, 10-19, 20-49, >49 (Supplementary Material Figure S7). The cytokine profile of a given disease is normalized by the corresponding bin the disease falls into: p-value = [number of diseases of which average cosine distances <given distance] / [total number of diseases in the bin], referred to as normalized average Association Scores (NAAS). We collected 171 well-studied diseases of which the literature sampling frequency of 79 cytokines are available in ImmuneXpresso for validation. We compared the predicted NAAS with the known literature sampling frequencies by calculating the Spearman correlation coefficients, where MatLab computes p-values for Spearman’s rank correlation coefficient using the exact permutation distributions.

### Analyze disease-specific cytokine profile features and subnetworks between pathogenesis and inflammation

Given a disease, we calculate the NAAS with each of the 126 cytokines, resulting a 126-dimension disease-specific cytokine profile. We analyzed the features of disease-specific cytokine profile via hierarchical clustering. The disease-specific cytokine profile is further converted into “Immune Scores” by averaging the NAAS of 47 inflammation related cytokines, 37 chemokines, 13 growth factors and 29 other cytokines, respectively, to suggest contribution from four aspects to inflammatory responses. In order to visualize the subnetwork formed between pathogenesis (disease associated genes) to inflammatory responses (essential cytokines), we label the disease genes with “disease-specific” and “cytokine receptors” by mapping to the 110 cytokine receptors that we defined by filtering genes downloaded from GeneCards ^32^. We graph the pairwise connections (NAAS > 0.8) between disease genes and cytokines into a force-directed layout that uses attractive forces between adjacent nodes and repulsive forces between distant nodes. To quantify the information exchange between pathogenesis and inflammatory responses, we calculated Immune Connection Density (ICD) by by 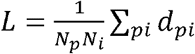, where *N_p_* is the number of total pathogenesis genes in the disease-specific network, *N_i_* is the number of total cytokines in the disease-specific network, and *d_pi_* is the cosine distance of the two sets of genes in embedding space^22^.

### Identify highly connected graphs between pathogenesis and inflammation by spectrum partition

For a given disease, we construct graph G using the predicted subnetworks derived from the high confidence interactions between pathogenesis genes, receptors and cytokines, with the interactions between cytokines removed. We calculate the Laplacian matrix L of the graph that is a square, symmetric, sparse matrix of G. The smallest non-null eigenvalue is called Fiedler value, which is the algebraic connectivity of a graph; the further from zero, the more connected. Fiedler vector w is the eigenvector corresponding to the smallest non-null eigenvalue of the graph. Partition the graph into two or three subgraphs using the Fiedler vector ***w***. A node is assigned to one subgraph if it has a positive value in ***w*** (well-connected nodes). Otherwise, the node is assigned to another subgraph (poorly connected nodes). Alternatively, the nodes of close to zero values in ***w*** can be placed in a class of their own (articulation point). This practice is called a sign cut or zero threshold cut. The sign cut minimizes the weight of the cut, subject to the upper and lower bounds on the weight of any nontrivial cut of the graph^33^.

## Notes

https://simtk.org/projects/cytokine

### Competing Interest Statement

The authors have declared no competing interest.

