## Supplemental Material for "Construction of Disease-specific Cytokine Profiles by Associating Disease Genes with Immune Responses"

Figure S1. We have inspected network sparsity within and between known functional modules in protein-protein interaction (PPI) networks. The connection rate is defined as the ratio of the number of actual links to the total number of links present in a complete graph of a given number of nodes. First, we found that the links within modules for immune response (Intra-ME) are under-represented, compared with the average connection rate within the heavily studied modules for metabolic, signaling. The low connection rates within the immune response modules result from the lack of informative resources about immune response. Second, we found that the associations between functional modules are not well defined, suggesting that we need to better define the associations between immune response and functional modules.

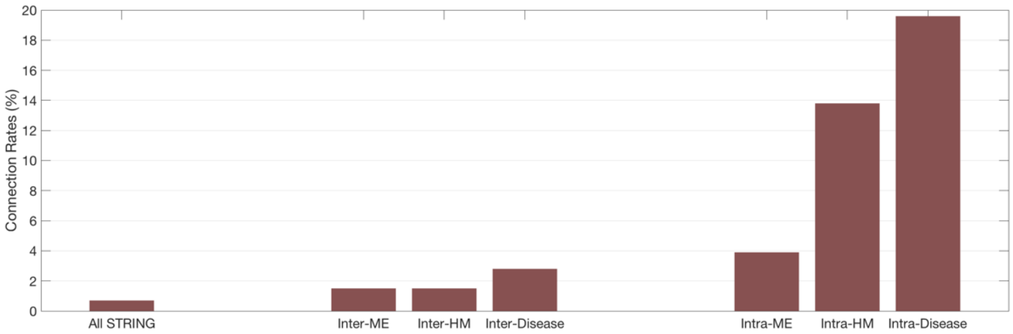

Figure S2. The predicted network distances of 8,521,944 edges correlate with the known confidence scores (low or medium confidence scores in STRING). The grey area shows the standard deviation of predicted distances of a give confidence score.

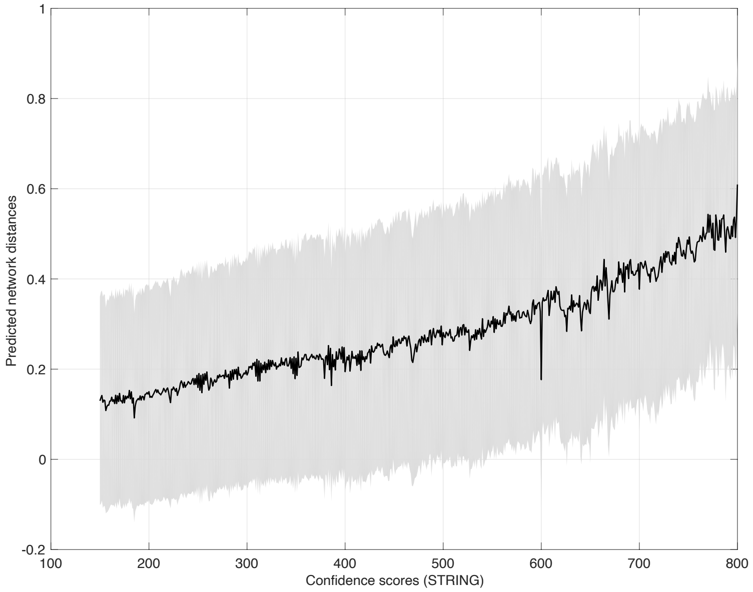

Figure S3. The distribution of the predicted network distances.
We calculated the probability of being associated of all possible pairs (108,140,571 pairs, shown in grey) between the 14,707 human genes. A total of 9,250,034 edges (shown in blue) are known with confidence scores in STRING (728,090 high confidence scores, and 9,250,034 low or medium confidence scores).

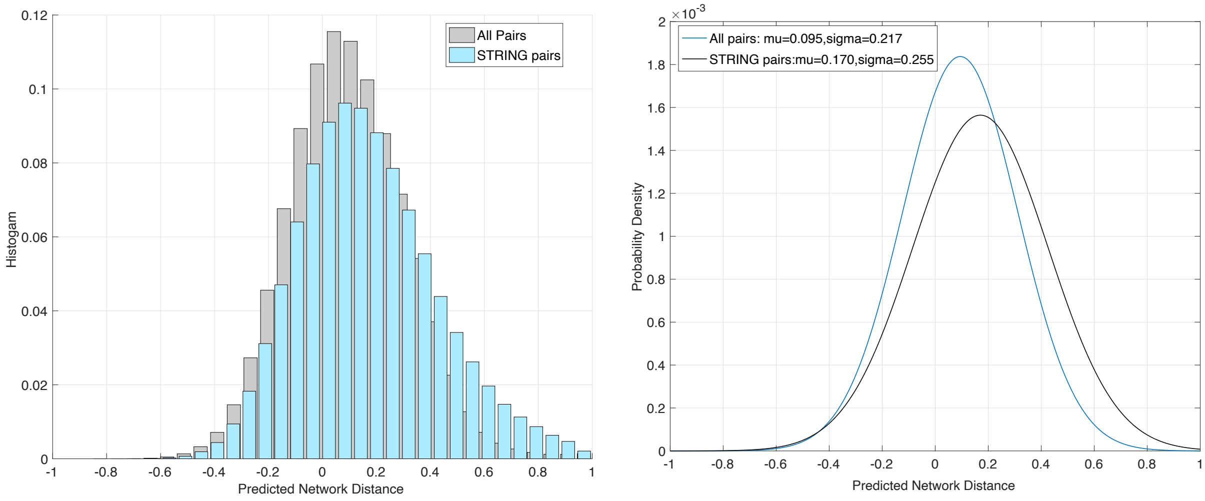

Figure S4. Predicted network distances correlate with the known confidence scores in STRING. A total of 9,250,034 edges are known with confidence scores (range from 100-1000). The differential box plots of predicted distances within each confidence bin suggest the predicted distances correlate with the known confidence in STRING.

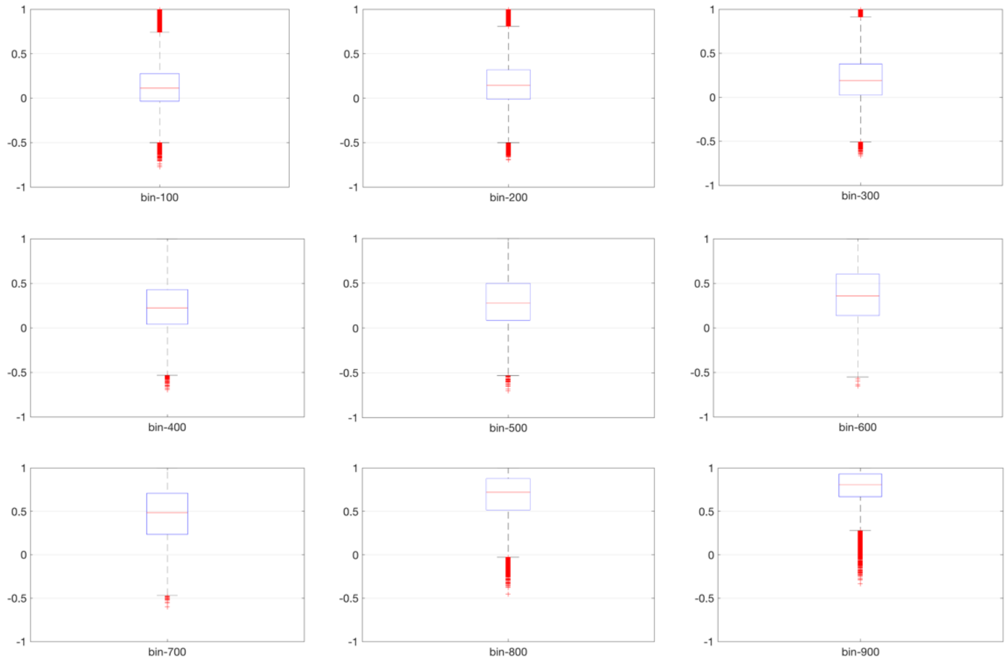

Figure S5. The histogram of the number of genes associated with each of the 171 diseases.

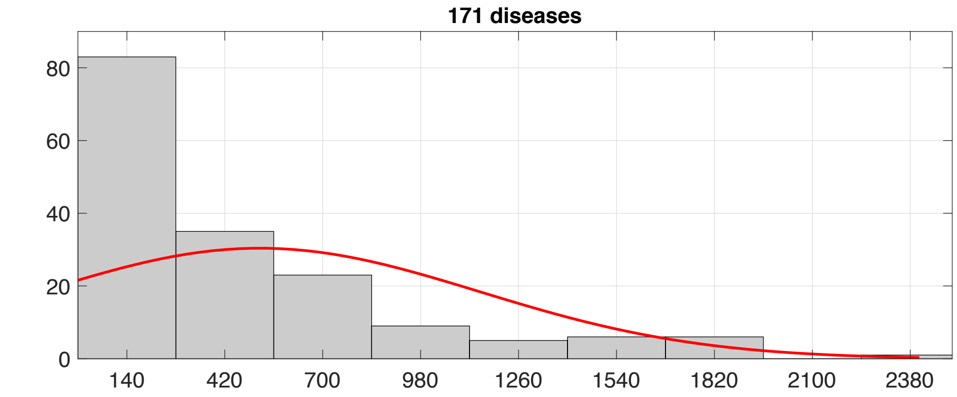

Figure S6. The relationship between the number of genes associated with each of the 171 diseases and the p-value that evaluates the correlation between predicted cytokine profiles and literature sampling frequency.

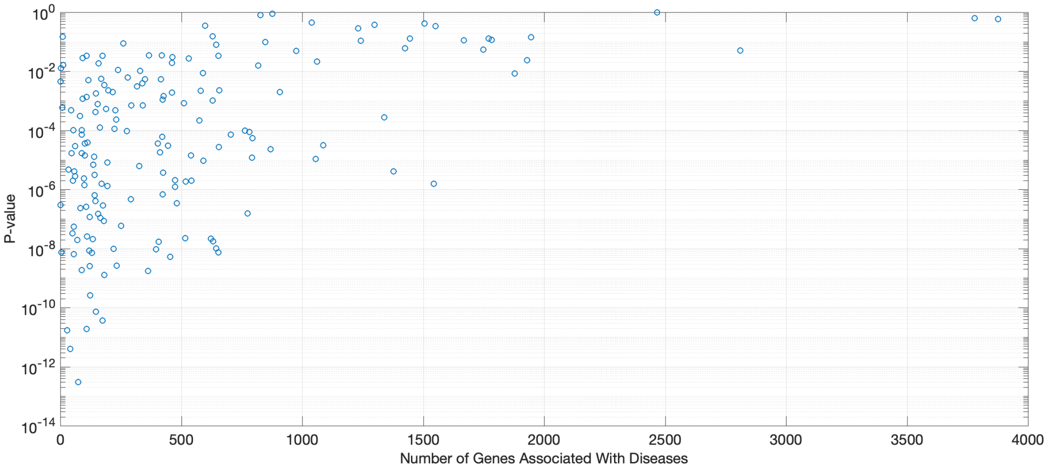

Figure S7 Bins defined by the number of genes associated with diseases. This was used for normalizations of cytokine profile of individual diseases.

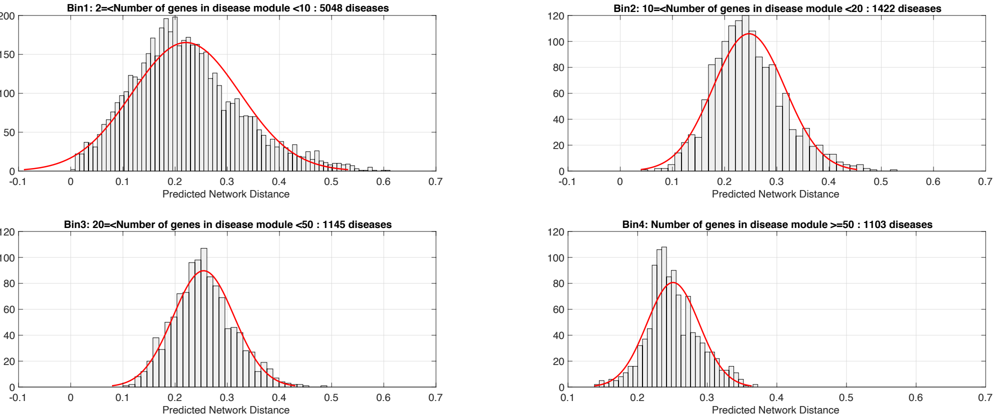

Figure S8. Graph plots showing interactions between pathogenesis genes (green and purple squares) to inflammatory responses (essential genes in dots). The graph was plotted in “force-directed layout that uses attractive forces between adjacent nodes and repulsive forces between distant nodes. For immune disorder SLE, a large number of pathogenesis genes (purple squares) are making interactions with core of inflammation cytokines (orange) and chemokines (dark red) directly or through a large number of receptors (green squares). As for infectious disease TB, fewer pathogenesis genes (purple squares) are making interactions with a large number of inflammation cytokines (orange) but are distant from chemokines (dark red). Attractive forces between pathogenesis and chemokine responses are observed in Metabolic Syndrome X, but not in Aneurysm or Acute Leukemia.

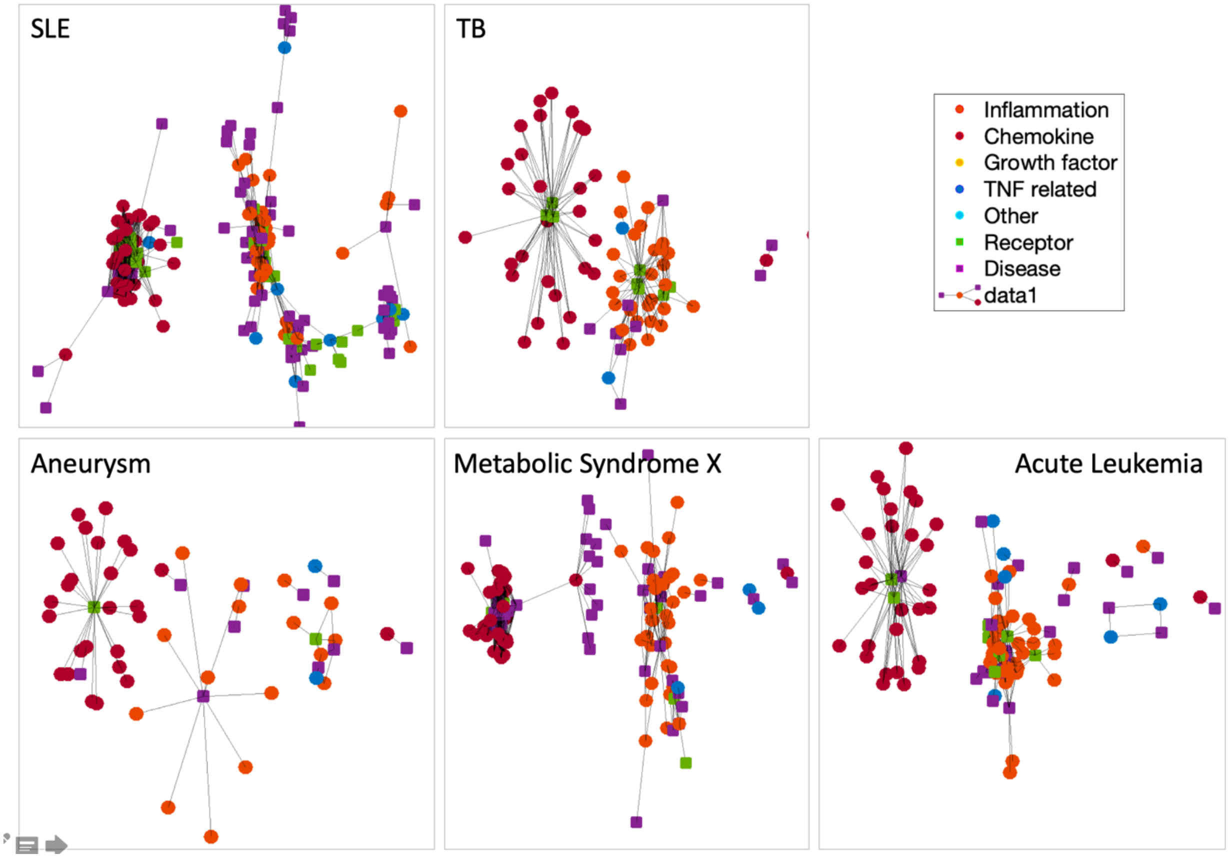

Figure S9. Graph showing information flow from pathogenesis genes (green and purple squares) to inflammatory responses (essential genes in dots) by plotting the high confidence interactions. The graph was plotted by placing genes into a set of layers, revealing their hierarchical structure.

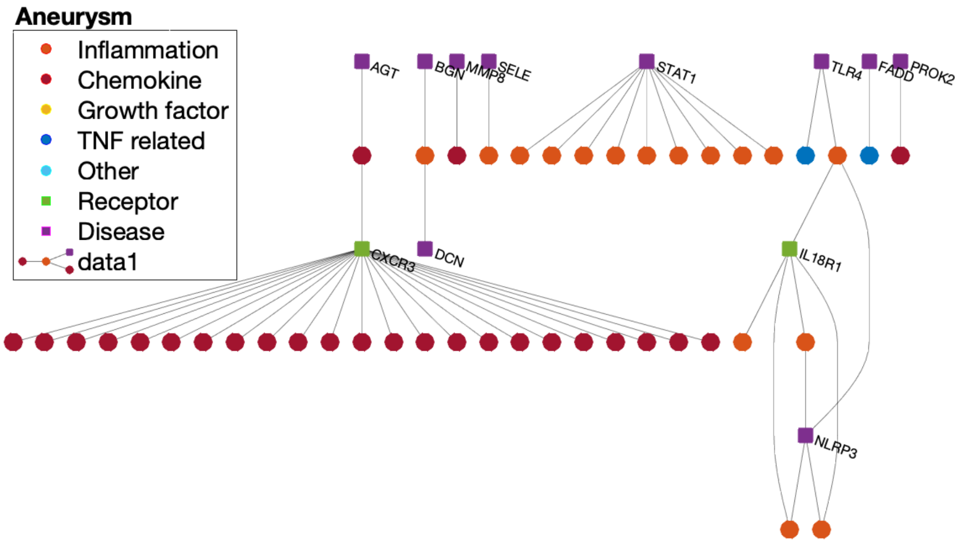

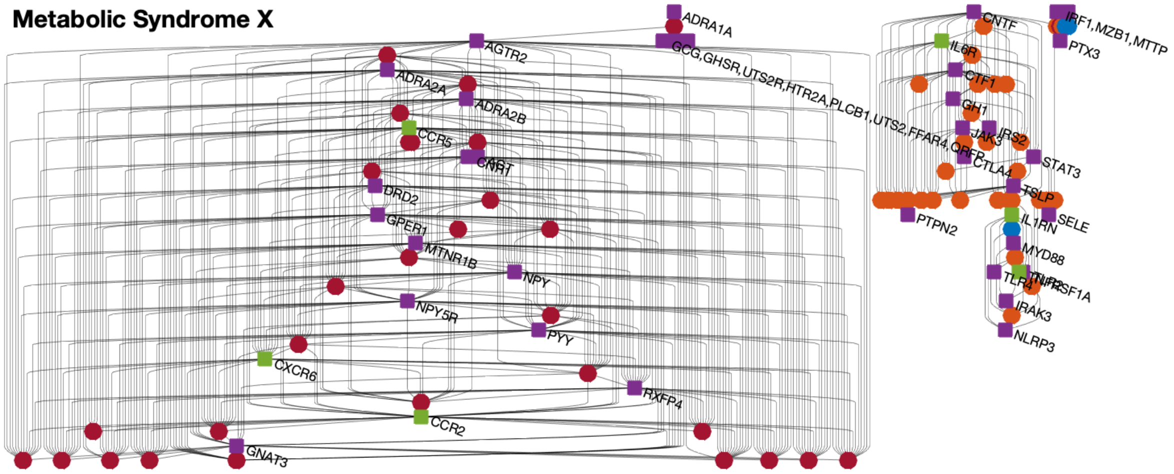

Table S1. Statistics of selected sets from STRING and three classes of functional modules. The fourth column 'Reference" calculates the number of possible connections, while the fifth column shows the observed connection rate.  When counting the connection observed in a gene set or between gene sets, we delete the genes observed in both sets.

| Subsets | # genes involved in edges with combined scores >0.8 | # Edges with combined scores >0.8 | Reference | Connection Rate |
|---|---|---|---|---|
| STRING-human | 14707 | 728090 | 108140571 | 0.67 |
| 47 sets of ME | 5657 | 261520 | 15997996 | 1.63 |
| Between ME | 5378 | 209064 | 14023305 | 1.49 |
| Within ME | 3360 | 29958 | 770775 | 3.89 |
| 50 sets of HM | 3805 | 131420 | 7237110 | 1.82 |
| Between HM | 2013 | 30018 | 2126204 | 1.41 |
| Within HM | 1181 | 7264 | 56801 | 12.79 |

Table S2. Gene signatures

| | |
|---|---|
| blue1 | RPS6, DOCK4, RPS9, RPS3A, RPS29, RPS28, RPS16, RPS21, RPS2, RPS3, RPS14, RPS23, RPS26, RPS18, RPS27L, RPS15, RPS11, MCTS1, RPS4X, RPS25, RPS19, RPS10, RPS24, RPS5, RPS17, RPS20, RPS7, FAU, RPS12, RPS13, RPS8, RPS15A, RPLP0, GSPT2, RPL8, RPL9, RPL32, RPL5, RPL12, RPL4, RPL13, RPL31, RPL18A, RPL7, RPL15, RPL30, RPL3, RPL6, RPL27, RPL11, RPL13A, RPL10A, RPL38, RPL26L1, RPL36, RPL35A, RPL10, RPL18, RPL28, RPL23, RPL37A, RPL17, RPL10L, RPL24, RPL19, RPL39, RPL26, RPL29, RPLP1, RPL3L, RPL7A, RPLP2, RPL39L, RPL22L1, RPL37, RSRC1, RPL22, RPL36A, RPL36AL, RPL21, RPL34, RPL14, RPL35, RPL27A, RPL23A, SRP54, C18orf32, RPL7L1, DIRC3, ETF1, SMG9, ABCF1, EIF3H, EIF4H, EIF5B, EIF3C, PELO, EIF3K, EIF3J, EIF3F, EIF3E, EIF3D, EIF1AX, EIF3M, EIF3L, EIF3CL, EEF1A2, SSR1, SSR3, SRPRB, TPT1, WDR31, SEC61A2, SEC11A, SEC61G, SRP72, ENSG00000215472, SRP14, SSR2, SRP68, SRP9, SPCS2, TRAM1, SERBP1, DENR, EIF5, EEF1D, ERI1, EIF3G, RPSAP58, SEC61B, SRP19, SRPR, SPCS3, SEC11C, SPCS1, NOP58, NOP56, BOP1, WDR12, PES1, RSL24D1, EIF6, HEATR1, PNO1, RRP9, DHX37, RPP30, RPP21, RPP14, RPP25, CIRH1A, BYSL, UTP3, NOL6, UTP14A, UTP11L, WDR75, DDX47, RRP36, RRP7A, EMG1, LTV1, RCL1, RIOK3, NOL11, UTP20, NOB1, FCF1, RIOK1, WBSCR22, UTP6, MPHOSPH10, WDR36, TBL3, UTP15, IMP4, PWP2, NOP14, KRR1, DDX52, NOC4L, BMS1, WDR43, UTP18, UTP14C, RIOK2, DDX49, DIEXF, WDR46, TSR1, WDR3, FBL, WDR18, EXOSC10, EXOSC3, EXOSC4, C1D, MPHOSPH6, NOL9, HBS1L, DIS3L2, DIS3, EXOSC1, EXOSC5, EXOSC8, EXOSC6, EXOSC7, EXOSC9, EXOSC2, PDCD11, MRTO4, IMP3, NSA2, RPP40, TTC37, ZBTB48, SUPV3L1, REXO4 |
| blue2 | KAT5, BTG1, RPA1, RPA2, RPA3, RFC5, RFC1, RFC2, RFC4, RFC3, POLE, POLE2, POLE3, ERCC1, ERCC4, POLE4, POLD3, POLD4, LIG1, MAD2L2, POLD1, DTL, CHRAC1, THUMPD1, WDR76, CHEK1, ATR, RAD51, RAD51C, BLM, TOP3A, BRCA2, EXO1, RAD51AP1, ATRIP, FANCD2, UBE2T, WDR48, USP1, FANCM, FANCE, FANCF, EME1, FANCG, POLN, FANCA, C19orf40, MUS81, SLX4, C17orf70, FANCB, FANCC, FANCL, EME2, C1orf86, DCLRE1B, DCLRE1A, FAN1, HELQ, WRN, RAD52, DNA2, XRCC3, XRCC2, RAD9B, RHNO1, SPIDR, RMI1, RAD51B, RAD17, RMI2, PALB2, TIMELESS, RAD1, HUS1, RAD9A, BRIP1, REV3L, RAD18, MSH2, MSH6, PIF1, MTMR12, |
| sig1 | APP, KNG1, PENK, C3, LPAR1, LPAR3, LPAR2, BDKRB1, CASR, MCHR2, NMUR1, LPAR5, PMCH, ANXA1, BDKRB2, NMU, NMUR2, NMS, GPR17, MCHR1, AGTR2, SAA1, C5, ADCY8, ADCY3, ADCY1, ADCY2, ADCY5, ADCY7, ADCY4, ADCY9, ADCY6, RLN3, GNAT3, SSTR3, GABBR2, HTR1B, GPR18, CCL28, HCAR3, NPW, TAS2R16, GRM8, RXFP3, NPY4R, NPY, SSTR2, CNR1, GRM6, GRM3, PTGDR2, APLN, SSTR4, ADORA3, P2RY13, TAS2R3, GPR31, TAS2R10, TAS2R7, CCR8, MTNR1B, OPRL1, S1PR5, RXFP4, TAS2R46, GPSM1, DRD3, GRM4, GPR183, HTR1A, CXCR5, TAS2R8, GPSM2, NPBWR1, HRH3, HTR5A, C5AR1, GRM7, GABBR1, SST, PNOC, TAS2R40, TAS1R1, TAS2R42, TAS1R3, CCR10, CCR9, SSTR1, TAS2R31, PYY, PDYN, TAS1R2, NPY1R, NPB, CCR4, TAS2R19, DRD4, APLNR, OPRK1, OXER1, OXGR1, CNR2, P2RY12, TAS2R4, NPBWR2, ADORA1, TAS2R1, TAS2R5, FPR3, TAS2R13, TAS2R60, INSL5, GPR55, NPY5R, HTR1E, SSTR5, HTR1F, TAS2R41, HTR1D, CORT, HCAR2, SUCNR1, TAS2R20, DRD2, GAL, TAS2R14, GPR37, PTGER3, CHRM4, TAS2R43, P2RY4, P2RY14, NPY2R, GALR1, HEBP1, GALR2, PCP2, GALR3, TAS2R39, TAS2R9, CXCR6, GRM2, TAS2R38, HRH4, TAS2R30, GPSM3, ACKR3, |

| | |
|---|---|
| | TAS2R50, CXCR2, OPRM1, HCAR1, CCR3, CXCR1, S1PR4, S1PR2, GPER1, S1PR3, ADRA2B, ADRA2C, CXCR3, CCL4L1, CX3CR1, MTNR1A, GPR37L1, OPRD1, PPY, CCR1, CCR2, ADRA2A, POMC, RGS13, RGS4, GPR61, PITPNM3, CXCR4, S1PR1, CCR5, CCR7, AGT, GNAI2, GNAI1, GNAI3, GPR78, RGS22, RGS5, PRKCD, EDN1, GNAZ, GNAO1, PRKCH, RGS1 |
| sig2 | STAT3, STAT6, STAT4, JAK1, TYK2, TBX21, STAT1, STAT5A, STAT5B, JAK3, IL13RA1, IL6R, IFNGR1, IFNGR2, SOCS3, SOCS1, PTPN1, PTPN2, KIT, IRS2, BCL6, JAK2, GH1, PRL, GH2, SOCS4, CSF2RA, CSF2RB, IL3RA, IL5RA, EPOR, SASH3, IKZF3, ENSG00000254469, IL2RG, IL2RB, IL2RA, NMI, STAT2, IFNA16, IFNA8, IFNA7, IFNA14, IFNA10, IFNA21, IFNA4, IFNA17, ENSG00000249624, IFNAR1, IFNE, IFNAR2, IFNW1, IL6ST, CNTF, LIFR, OSMR, IL4R, GHR, CRLF1, PRLR, CSH1, CTF1, PTPRE, MPL, IL15RA, IL10RB, IL22RA1, IL20RA, IL20RB, IL10RA, IL27RA, CNTFR, CRLF2, IL21R, IFNLR1, IL22RA2, IL11RA, CSH2, IL31RA, C19orf60, IL9R, CSF3R, IL12RB1, IL23R, IL12RB2, TSLP, IL13RA2, IRF9, USP41, IKZF1, KLRD1, KIR2DL3, KLRC1, KIR2DL1, KIR3DL1, LAG3, CD207, MRC2, ENSG00000255819, KLRK1, HCST, NCR1, NCR3, NCR2, KLRC4, IFNA5, SFTPC, SFTPB, SFTPD, SFTPA2, SFTA3, SFTPA1, FOXP4, PITPNA, FOXP2, GNLY, NKG7, PTPN5, FCER2, BATF, GLG1, GRAP, GAB4, FCRLA, FAM132B, TIE1, PTPRU, FLT3, STAP1, ALK, SH2B3, EML4, TFF2, DOK6, ANGPT2, ANGPT4, BTLA, VTCN1, DIRAS2, S100A13, LMO4, EOMES, PRF1, SP110, SPN, CR2, IKZF4, SOCS7, ICAM1, CD1B, CD274, CD28, CTLA4, SELL, CD19, CD34, CD5, MRC1, CD1E, CD1D, CD83, TNFRSF18, TNFSF18, ICOS, B7RP1, PTPRCAP, ICOSL, IL17RC, CLEC7A, FOXP1, CLEC1A, CD276, STAP2, HLX, RETNLB, IFNA13, TLR4, MYD88, IRAK4, IRAK1, TLR1, TLR2, IRAK2, IL1RAP, TLR9, TLR3, TIRAP, MALT1, IRAK3, TXNIP, MAP3K1, FASLG, MAP3K5, TNFRSF14, CD40, TNFRSF9, TLR7, TLR8, CNPY3, UNC93B1, TLR5, S100A8, IL1R2, IL1RL1, NRK, TMEM26, S100A9, LY86, FOXP3, CD27, CD69, SELE, IL1R1, IL18R1, IL18RAP, SIGIRR, IL37, IL18BP, LBP, KLF2, CCL3L3, GPR29, TNFRSF4, IDO1 |
| sig3 | UBA52, RPS27A, UBC, UBB, IFIT1B, DDI1, LYST, TMEM106B, TULP2, CTNNB1, YWHAE, YWHAG, YWHAB, PPP2CB, PPP2R1B, PPP2R5A, PPP2R5C, PPP2R5D, PPP2R5E, PPP2R5B, GSK3B, AXIN1, APC, AJUBA, CSNK1G2, CSNK1G1, FRAT2, FRAT1, GSK3A, APC2, MEA1, PMF1-BGLAP, COL4A3BP, NINJ1, EIF2AK4, C1orf111, DVL2, CFTR, CCDC88C, UBE2D1, ITCH, SMURF2, WWP1, SKP1, CUL1, SKP2, UBE2E1, CUL3, VHL, CUL2, FBXL7, ANAPC4, CDC23, ANAPC10, CDC26, ANAPC5, ANAPC2, CDC27, ANAPC7, ANAPC11, ANAPC1, FZR1, BTRC, UBE2C, CDC16, EPAS1, PSMB8, ANAPC15, ANAPC16, HIF3A, EGLN3, EGLN1, HIVEP3, PSMD1, PSMC4, PSMC5, PSMC2, PSMC1, CSNK1A1, AMER1, AXIN2, PSMA4, PSMC3, SHFM1, PSMD4, PSMD3, PSMD14, PSMB5, PSMA3, PSMA5, PSMA2, PSMB2, PSMD2, PSMA7, PSMD11, PSMD12, PSMA1, PSMB3, PSMD7, PSME3, PSMA6, PSMB4, PSMB9, PSMB11, PSMD13, PSMF1, PSMB1, PSMD5, PSME4, PSMD6, PSMB6, PSMD10, PSMD9, PSMC6, PSME1, PSMA8, PSMB10, PSMD8, PSMB7, PSME2, TP73, TNKS, TNKS2, MAPK6, PTGES3L, WTIP, LIMD1, SPOPL, PRICKLE1, ADRM1, RNF146, ZSWIM8, HECW1, UBD, PSMG2, CCDC74B, SPOP, OAZ2, POMP, KIAA2012, SMIM15, USP14, CCDC92, GNB2L1, HABP4, CASP4, EIF2AK1, GLI2, SUFU, GLI3, GLI1, IHH, DHH, CDON, PTCH1, BOC, GAS1, PTCH2, PAK2, NUMB, NUMBL, SHH, RUNX3, DISP1, AURKA, FBXO5, PTTG1, GTSE1, UCHL5, DZIP1, MYSM1, PARP6, KLHL12, ASNA1, SGTA, IGBP1, ENSA, CCNG2, PPP2R3C, SENP2, KLHDC3, QTRT1, QTRTD1, TXNL1, USP37, TBC1D20, USP6 |

| | |
|---|---|
| sig4 | APOB, TF, LAMB1, LAMC1, LAMB2, ADAM10, DNAJC3, FUCA2, P4HB, CTAGE6, HSP90B1, CDH2, QSOX1, CST3, HP, ORM2, APCS, SERPINA7, CKAP4, AHSG, GAS6, SERPINC1, IGFBP1, MSLN, MFI2, APOA5, PDIA6, MBTPS1, PNPLA2, CP, C4A, PRKCSH, TNC, CALU, IGFBP4, SPP2, CHRDL1, APOL1, PRSS23, EVA1A, FSTL3, IGFBP7, SCG3, MFGE8, AFP, STC2, DMP1, WFS1, VGF, FAM20C, APLP2, VWA1, AMTN, LTBP1, GOLM1, KTN1, MIA3, AMBN, HRC, TMEM132A, LGALS1, FSTL1, BPIFB2, ENAM, MEPE, MATN3, ITIH2, ANO8, SPARCL1, MXRA8, SERPIND1, SHISA5, SERPINA10, AMELX, SCG2, MGAT4A, FAM20A, CYR61, FBN1, IGFBP5, BMP15, NUCB1, PCSK9, NOTUM, CHGB, RCN1, PROC, IGFBP3, KLK4, APOA1, APOA2, GPC3, APOE, VCAN, MSR1, F2, CPB2, PROCR, SERPINA1, F5, F8, PAPPA2, TYRO3, VTI1B, THBS1, CSRP1, GP1BA, VTN, F10, SERPINA5, F9, F13B, GP5, GP9, GP1BB, F11, F12, KLKB1, SERPINE2, RIC8B, FN1, FGG, FGA, VWF, FGB, ACTN2, ACTN1, ACTN4, ALB, IGF2, KLK1, SERPINE1, A2M, CLU, SPARC, FERMT3, TMSB4X, PROS1, OLA1, F13A1, TEX264, SERPINF2, FAM49B, HRG, NHLRC2, SCCPDH, MMRN1, CFD, SRGN, MAGED2, GTPBP2, TOR4A, PCYOX1L, APOOL, LEFTY2, SERPING1, ISLR, DNAJC4, ALDOA, ORM1, PLG, FIGF, VEGFB, VEGFC, A1BG, TIMP1, REN, ACE, ENSG00000264813 |
| sig5 | GNB1, GNGT1, FPR2, GNG2, GNG13, GNG8, GNG11, GNG12, GNG7, GNB2, GNG4, GNGT2, GNB4, GNB3, GNB5, GNG10, GNG5, GNG3, CHRM2, GRK6, GNAL, C3AR1, PSAP, FPR1, PCSK1N, GPRC5C, TRPM5, RLN1, RTP4, OR13G1, ADORA2B, PTH, SLC5A1, KIAA1919, SLC15A1, RGS2, RGS21, GHRL, AGRP, PTAFR, ARR3, NLN, ARHGEF1, ARHGEF12, GNA12, CYTH4, ARHGEF5, AKAP13, VR1, AKAP1, GCG, NPS, P2RY11, GPR39, GCGR, GNAS, GNAQ, PLCB2, TBXA2R, GNA11, GNA14, EDNRA, GNA15, OXT, PLCB1, P2RY2, PTGER1, PLCB3, PLCB4, GNRH1, GNRH2, GPR132, AVPR1B, HCRT, CHRM5, EDNRB, XCR1, HCRTR1, QRFPR, CHRM3, AVPR1A, GHSR, F2RL3, OXTR, HTR2C, P2RY10, MLN, KISS1R, CCK, P2RY6, GPR143, TAC1, NPFFR2, KISS1, LPAR6, NMBR, GRPR, NMB, HTR2A, TACR2, LTB4R2, HCRTR2, FFAR4, NTSR1, PROK1, ADRA1B, PROKR2, UTS2R, GNRHR, GPR65, CYSLTR1, EDN3, F2RL1, GPRC6A, CYSLTR2, GRP, ADRA1D, QRFP, XCL2, MLNR, UTS2B, EDN2, GPR68, ADRA1A, CCKBR, LTB4R, PTGFR, CCKAR, FFAR2, BRS3, NPFF, NTS, FFAR3, TACR3, LPAR4, TRH, PROK2, UTS2, NTSR2, TRHR, GAST, HTR2B, NPFFR1, TAC3, ARHGEF25, P2RY1, HRH1, FFAR1, GPR4, OPN4, GRK5, GRM1, GRM5, CHRM1, F2R, F2RL2, ADRBK1, TRIO, KALRN, GNA13, KCNJ3, S100A11, CD68, ACE2, SPG7, PXYLP1, GALP, C9, C8B, C8A, C6, C8G, C7, PPIF, CHID1, CHI3L2, STAB1, ITIH4, CR1 |
| sig6 | TRAF2, BIRC3, BIRC2, NFKBIB, REL, RELB, MAP3K14, NFKBIA, NFKB2, TRAF3, TRAF6, TAB3, RIPK1, BCL10, TNFRSF1A, CARD11, RIPK2, NOD2, TICAM2, TICAM1, FADD, TRADD, TNFRSF25, FAS, RIPK3, IKBKG, IKBKB, CHUK, MAP2K4, MAP2K7, MAP3K3, MAP4K4, CRADD, TRAF1, TRAF5, TNFRSF10B, NSMAF, TNIK, BCL3, TLR6, PELI2, PELI1, PELI3, MAP3K1, FASLG, MAP3K5, TLR3, MALT1, TIRAP, TXNIP, NLRP3, SETD6, SETD4, NKIRAS2, RALGAPA1, GFOD2, GFOD1, DDX58, IFIH1, EIF2AK2, TBK1, MAVS, IKBKE, ZBP1, TANK, RNF135, AZI2, CMC1, NLRX1, BID, NOD1, MAP3K8, TNFAIP3, OTUD7B, USP2, CYLD, USP4, RNF31, SHARPIN, TAX1BP1, OTUB2, MADD, MLKL, PAWR, CARD10, CARD14, TNFRSF10A, TNFSF10, CFLAR, MAP4K2, TNFRSF17, PIDD1, TNFRSF10C, TNFRSF10D, TNIP1, IFI16, TBKBP1, ENSG00000026036, TNFRSF6B, TNFRSF13B, MAP3K7, USP21, TAB1, TAB2, XIAP, ERN1, CD14, LY96, TNIP2, PTPN11, GAB2, IRS2, |

| | |
|---|---|
| | PTPN1, PTPN2, KIT, TEK, ANGPT1, CSF1R, PLCG1, GRB2, SHC1, CRKL, PDGFRA, GAB1, IGF1R, PIK3CB, DIRAS1, PIK3CD, GDNF, IRS1, IRS4, BDNF, PRRT2, ERBB2, HBEGF, ERBB4, FLT1, SHC2, PTPN12, PGF, INSRR, FLT4, FGF13, CBLB, FCER1G, VAV2, VAV3, IGHV3-11, IGLL5, C6orf25, LCP2, ZAP70, GRAP2, FYB, SLA2, BLNK, PIK3AP1, DAPP1, SLA, IGHV4-38-2, NCAM1, ST8SIA4, LAT, UBASH3A, SYK, BTK, ITK, TRAT1, CD8A, CD8B, PAG1, BLK, PLCG2, CD79A, MS4A2, CD79B, FCGR2B, INPP5D, FCER1A, POU2AF1, FER, PDPN, INPPL1, BAG4, FES, PTPN13, BMX, DOK3, BCR, SOS2, NTF3, AXL, NTRK3, RIT1, SHC3, GRB7, NTF4, RIT2, RET, GFRA1, GFRA3, ARTN, PSPN, GFRA2, GFRA4, RAP1GAP, NRTN, DOK5, DOK1, SH2B2, DOK2, SHC4, GRB10, FLT3LG, KL, FGFR3, ZMYM2, MYO18A, FGF21, LRPPRC, TFF3, TNK2, TYROBP, TREM2, ENSG00000255641, TEC, SH3BP5, IBTK, CD22, KLRG1, CD72, CLEC1B, CD300LB, LAT2, TXK, IGLL1, VPREB3, TRPV6, SH2D2A, SLAMF1, CLEC6A, NIPSNAP1, UNC5C, UNC5D, PLEKHA2, RASGRP3, S100G, DOK4, LZTR1, CADPS2, LY6G6F, CD2, CD7, LGALS9, MME, CD48, MAP4K1, SH3BP2, CLEC4E |

Table S3. The predicted cytokine profiles correlate with the known literature sampling frequency in ImmuneXpresso for the 171 well-studied diseases.

| Disease ID | #gene | p-value | CC | Disease name | Category | Category |
|---|---|---|---|---|---|---|
| C1290886 | 41 | 8.78E-12 | 0.67525 | Chronic inflammatory disorder | C23 | "Pathological Conditions, Signs and Symptoms" |
| C0023290 | 74 | 9.39E-12 | 0.67454 | "Leishmaniasis, Visceral" | C01 | Infections |
| C0029118 | 29 | 1.39E-11 | 0.67044 | Opportunistic Infections | C01 | Infections |
| C1290884 | 174 | 3.88E-11 | 0.65944 | Inflammatory disorder | C23 | "Pathological Conditions, Signs and Symptoms" |
| C0275524 | 147 | 7.29E-11 | 0.65244 | Coinfection | C01 | Infections |
| C0026946 | 123 | 2.71E-10 | 0.63728 | Mycoses | C01 | Infections |
| C0035235 | 109 | 8.77E-10 | 0.62294 | Respiratory Syncytial Virus Infections | C01 | Infections |
| C0004623 | 234 | 1.85E-09 | 0.61346 | Bacterial Infections | C01 | Infections |
| C0011615 | 362 | 2.73E-09 | 0.60837 | Dermatitis, Atopic | C16;C17;C20 | "Congenital, Hereditary, and Neonatal Diseases and Abnormalities; Skin and Connective Tissue Diseases; Immune System Diseases" |
| C0243026 | 454 | 5.56E-09 | 0.59883 | Sepsis | C23;C01 | "Pathological Conditions, Signs and Symptoms; Infections" |
| C0009324 | 654 | 7.26E-09 | 0.59518 | Ulcerative Colitis | C06 | Digestive System Diseases |
| C0151317 | 5 | 7.59E-09 | 0.59457 | Chronic infection | C01;C20 | Infections; Immune System Diseases; |

| | | | | | | |
|---|---|---|---|---|---|---|
| C0024530 | 396 | 9.74E-09 | 0.5911 | Malaria | C01 | Infections |
| C0021390 | 645 | 1.03E-08 | 0.59026 | Inflammatory Bowel Diseases | C06 | Digestive System Diseases |
| C0007570 | 220 | 1.25E-08 | 0.58761 | Celiac Disease | C06;C18 | Digestive System Diseases; Nutritional and Metabolic Diseases |
| C0036690 | 406 | 1.71E-08 | 0.58316 | Septicemia | C23;C01 | "Pathological Conditions, Signs and Symptoms; Infections" |
| C0019693 | 631 | 2.03E-08 | 0.58067 | HIV Infections | C01;C20 | Infections; Immune System Diseases |
| C0010346 | 622 | 2.30E-08 | 0.57884 | Crohn Disease | C06 | Digestive System Diseases |
| C0009319 | 516 | 2.31E-08 | 0.57878 | Colitis | C06 | Digestive System Diseases |
| C0155877 | 181 | 2.47E-08 | 0.57779 | Allergic asthma | C08;C20 | Respiratory Tract Diseases; Immune System Diseases |
| C0876973 | 70 | 3.38E-08 | 0.57319 | Infectious Lung Disorder | C01;C08 | Infections; Respiratory Tract Diseases |
| C2607914 | 131 | 3.38E-08 | 0.57316 | Allergic rhinitis (disorder) | C08;C20;C09 | Respiratory Tract Diseases; Immune System Diseases; Otorhinolaryngologic Diseases |
| C0242966 | 56 | 4.52E-08 | 0.56884 | Systemic Inflammatory Response Syndrome | C23 | "Pathological Conditions, Signs and Symptoms" |
| C0023281 | 56 | 5.36E-08 | 0.56624 | Leishmaniasis | C01;C17 | Infections; Skin and Connective Tissue Diseases |
| C0024131 | 252 | 8.27E-08 | 0.55958 | Lupus Vulgaris | C01;C17 | Infections; Skin and Connective Tissue Diseases |

| | | | | | | |
|---|---|---|---|---|---|---|
| C0040100 | 180 | 9.23E-08 | 0.55789 | Thymoma | C04;C15 | Neoplasms; Hemic and Lymphatic Diseases |
| C0266929 | 119 | 1.06E-07 | 0.55569 | Chronic Periodontitis | C07 | Stomatognathic Diseases |
| C0025007 | 121 | 1.22E-07 | 0.55353 | Measles | C01 | Infections |
| C0026936 | 50 | 1.25E-07 | 0.55311 | Mycoplasma Infections | C01 | Infections |
| C0026918 | 88 | 1.27E-07 | 0.55281 | Mycobacterium Infections | C01 | Infections |
| C0036202 | 176 | 1.43E-07 | 0.55099 | Sarcoidosis | C15 | Hemic and Lymphatic Diseases |
| C0041327 | 135 | 1.51E-07 | 0.55013 | "Tuberculosis, Pulmonary" | C01;C08 | Infections; Respiratory Tract Diseases |
| C0042769 | 774 | 1.54E-07 | 0.54974 | Virus Diseases | C01 | Infections |
| C0014038 | 107 | 2.28E-07 | 0.54343 | Encephalitis | C10 | Nervous System Diseases |
| C0031099 | 157 | 2.36E-07 | 0.54288 | Periodontitis | C07 | Stomatognathic Diseases |
| C0031154 | 110 | 3.32E-07 | 0.53723 | Peritonitis | C06;C01 | Digestive System Diseases; Infections |
| C0041296 | 482 | 3.35E-07 | 0.53705 | Tuberculosis | C01 | Infections |
| C0011311 | 141 | 4.11E-07 | 0.53363 | Dengue Fever | C01 | Infections |
| C0023343 | 121 | 4.49E-07 | 0.5321 | Leprosy | C01 | Infections |
| C3714636 | 292 | 4.76E-07 | 0.53113 | Pneumonitis | C01;C08 | Infections; Respiratory Tract Diseases |
| C0023470 | 423 | 6.64E-07 | 0.52539 | Myeloid Leukemia | C04 | Neoplasms |
| C0019829 | 475 | 1.21E-06 | 0.51474 | Hodgkin Disease | C04;C20;C15 | Neoplasms; Immune System Diseases; Hemic and Lymphatic Diseases |

| | | | | | | |
|---|---|---|---|---|---|---|
| C0003872 | 171 | 1.41E-06 | 0.51206 | "Arthritis, Psoriatic" | C17;C05 | Skin and Connective Tissue Diseases; Musculoskeletal Diseases |
| C0023418 | 1544 | 1.44E-06 | 0.51165 | leukemia | C04 | Neoplasms |
| C0021400 | 474 | 1.76E-06 | 0.50804 | Influenza | C01;C08 | Infections; Respiratory Tract Diseases |
| C0003864 | 518 | 1.94E-06 | 0.50621 | Arthritis | C05 | Musculoskeletal Diseases |
| C0033860 | 542 | 2.08E-06 | 0.50492 | Psoriasis | C17 | Skin and Connective Tissue Diseases |
| C0038013 | 194 | 2.21E-06 | 0.50382 | Ankylosing spondylitis | C05 | Musculoskeletal Diseases |
| C1264606 | 62 | 2.96E-06 | 0.49836 | Persistent infection | C23;C01 | "Pathological Conditions, Signs and Symptoms; Infections" |
| C0017152 | 141 | 3.05E-06 | 0.49774 | Gastritis | C06 | Digestive System Diseases |
| C0085669 | 425 | 3.61E-06 | 0.49454 | Acute leukemia | C23;C04 | "Pathological Conditions, Signs and Symptoms; Neoplasms" |
| C0042164 | 84 | 4.46E-06 | 0.4905 | Uveitis | C11 | Eye Diseases |
| C0023467 | 1378 | 4.49E-06 | 0.49037 | "Leukemia, Myelocytic, Acute" | C04 | Neoplasms |
| C0025289 | 53 | 5.05E-06 | 0.48805 | Meningitis | C10 | Nervous System Diseases |
| C0240066 | 58 | 5.08E-06 | 0.48795 | Iron deficiency | C18 | Nutritional and Metabolic Diseases |
| C0042384 | 97 | 5.78E-06 | 0.48543 | Vasculitis | C14 | Cardiovascular Diseases |
| C0011603 | 165 | 6.04E-06 | 0.48455 | Dermatitis | C17 | Skin and Connective Tissue Diseases |

| | | | | | | |
|---|---|---|---|---|---|---|
| C13067 59 | 145 | 6.14E-06 | 0.48423 | Eosinophilic disorder | C15 | Hemic and Lymphatic Diseases |
| C00064 13 | 326 | 6.30E-06 | 0.48371 | Burkitt Lymphoma | C04;C01;C20; C15 | Neoplasms; Infections; Immune System Diseases; Hemic and Lymphatic Diseases |
| C00361 17 | 34 | 6.69E-06 | 0.48254 | Salmonella infections | C01 | Infections |
| C00029 40 | 136 | 6.82E-06 | 0.48214 | Aneurysm | C14 | Cardiovascular Diseases |
| C00267 64 | 105 6 | 9.31E-06 | 0.4759 | Multiple Myeloma | C04;C20;C15; C14 | Neoplasms; Immune System Diseases; Hemic and Lymphatic Diseases; Cardiovascular Diseases |
| C00797 31 | 592 | 9.65E-06 | 0.47515 | B-Cell Lymphomas | C04;C20;C15 | Neoplasms; Immune System Diseases; Hemic and Lymphatic Diseases |
| C00241 41 | 793 | 1.02E-05 | 0.474 | "Lupus Erythematosus, Systemic" | C17;C20 | Skin and Connective Tissue Diseases; Immune System Diseases |
| C00011 75 | 140 | 1.13E-05 | 0.47198 | Acquired Immunodeficienc y Syndrome | C01;C20 | Infections; Immune System Diseases |
| C00270 59 | 99 | 1.19E-05 | 0.47085 | Myocarditis | C14 | Cardiovascular Diseases |
| C00083 54 | 101 | 1.39E-05 | 0.46761 | Cholera | C01 | Infections |
| C00061 18 | 541 | 1.46E-05 | 0.46665 | Brain Neoplasms | C04;C10 | Neoplasms; Nervous System Diseases |
| C00352 43 | 47 | 1.48E-05 | 0.46628 | Respiratory Tract Infections | C01;C08 | Infections; Respiratory Tract Diseases |

| | | | | | | |
|---|---|---|---|---|---|---|
| C0376545 | 413 | 1.79E-05 | 0.46237 | Hematologic Neoplasms | C04;C15 | Neoplasms; Hemic and Lymphatic Diseases |
| C0023434 | 870 | 2.30E-05 | 0.45696 | Chronic Lymphocytic Leukemia | C04;C20;C15 | Neoplasms; Immune System Diseases; Hemic and Lymphatic Diseases |
| C0524909 | 194 | 2.31E-05 | 0.45686 | "Hepatitis B, Chronic" | C06;C01 | Digestive System Diseases; Infections |
| C0024305 | 445 | 2.79E-05 | 0.45277 | "Lymphoma, Non-Hodgkin" | C04;C20;C15 | Neoplasms; Immune System Diseases; Hemic and Lymphatic Diseases |
| C0024299 | 1086 | 3.11E-05 | 0.45043 | Lymphoma | C04;C20;C15 | Neoplasms; Immune System Diseases; Hemic and Lymphatic Diseases |
| C0019163 | 656 | 3.16E-05 | 0.45008 | Hepatitis B | C06;C01 | Digestive System Diseases; Infections |
| C0023487 | 403 | 3.38E-05 | 0.44861 | Acute Promyelocytic Leukemia | C04 | Neoplasms |
| C0266999 | 112 | 3.76E-05 | 0.44624 | Vesicular Stomatitis | C01;C07;C22 | Infections; Stomatognathic Diseases; Animal Diseases |
| C0242497 | 1 | 3.98E-05 | 0.44497 | Intestinal schistosomiasis | C01 | Infections |
| C0026769 | 795 | 5.20E-05 | 0.43891 | Multiple Sclerosis | C20;C10 | Immune System Diseases; Nervous System Diseases |
| C0019348 | 422 | 6.26E-05 | 0.43466 | Herpes Simplex Infections | C01;C17 | Infections; Skin and Connective Tissue Diseases |
| C0039103 | 88 | 6.55E-05 | 0.43361 | Synovitis | C05 | Musculoskeletal Diseases |

| | | | | | | |
|---|---|---|---|---|---|---|
| C0011854 | 705 | 7.05E-05 | 0.43191 | Diabetes Mellitus, Insulin-Dependent | C18;C20;C19 | Nutritional and Metabolic Diseases; Immune System Diseases; Endocrine System Diseases |
| C0149516 | 62 | 8.45E-05 | 0.42768 | Chronic sinusitis | C01;C08;C09 | Infections; Respiratory Tract Diseases; Otorhinolaryngologic Diseases |
| C0001339 | 88 | 8.74E-05 | 0.42687 | Acute pancreatitis | C06 | Digestive System Diseases |
| C0026948 | 102 | 9.01E-05 | 0.42615 | Mycosis Fungoides | C04;C20;C15 | Neoplasms; Immune System Diseases; Hemic and Lymphatic Diseases |
| C0023449 | 781 | 9.06E-05 | 0.42603 | Acute lymphocytic leukemia | C04;C20;C15 | Neoplasms; Immune System Diseases; Hemic and Lymphatic Diseases |
| C0018213 | 275 | 9.67E-05 | 0.42446 | Graves Disease | C11;C20;C19 | Eye Diseases; Immune System Diseases; Endocrine System Diseases |
| C0023473 | 764 | 0.00010036 | 0.42359 | "Myeloid Leukemia, Chronic" | C04;C15 | Neoplasms; Hemic and Lymphatic Diseases |
| C0036323 | 54 | 0.00010827 | 0.42177 | Schistosomiasis | C01 | Infections |
| C0019159 | 224 | 0.00011682 | 0.41994 | Hepatitis A | C06;C01 | Digestive System Diseases; Infections |
| C0023443 | 89 | 0.00014729 | 0.41429 | Hairy Cell Leukemia | C04;C20;C15 | Neoplasms; Immune System Diseases; Hemic and Lymphatic Diseases |

| | | | | | | |
|---|---|---|---|---|---|---|
| C0024143 | 164 | 0.00017734 | 0.40968 | Lupus Nephritis | C13;C17;C12;C20 | Female Urogenital Diseases and Pregnancy Complications; Skin and Connective Tissue Diseases; Male Urogenital Diseases; Immune System Diseases |
| C3463824 | 575 | 0.00021196 | 0.4052 | MYELODYSPLASTIC SYNDROME | C15 | Hemic and Lymphatic Diseases |
| C0010823 | 232 | 0.00022847 | 0.4033 | Cytomegalovirus Infections | C01 | Infections |
| C0003873 | 1340 | 0.00027826 | 0.39823 | Rheumatoid Arthritis | C17;C05;C20 | Skin and Connective Tissue Diseases; Musculoskeletal Diseases; Immune System Diseases |
| C0001486 | 82 | 0.00028324 | 0.39778 | Adenovirus Infections | C01 | Infections |
| C0018799 | 228 | 0.00048032 | 0.38378 | Heart Diseases | C14 | Cardiovascular Diseases |
| C0024266 | 45 | 0.00052347 | 0.38145 | Lymphocytic Choriomeningitis | C01;C10 | Infections; Nervous System Diseases |
| C0023440 | 190 | 0.00053316 | 0.38094 | Acute Erythroblastic Leukemia | C04;C15 | Neoplasms; Hemic and Lymphatic Diseases |
| C3812396 | 8 | 0.00060645 | 0.37741 | Chronic idiopathic pulmonary fibrosis | C08 | Respiratory Tract Diseases |
| C0017658 | 145 | 0.00066619 | 0.3748 | Glomerulonephritis | C13;C12 | Female Urogenital Diseases and Pregnancy |

| | | | | | | |
|---|---|---|---|---|---|---|
| | | | | | | Complications; Male Urogenital Diseases |
| C0153690 | 293 | 0.00067075 | 0.37461 | Secondary malignant neoplasm of bone | C23;C04;C05 | "Pathological Conditions, Signs and Symptoms; Neoplasms; Musculoskeletal Diseases" |
| C0740457 | 341 | 0.00074817 | 0.37155 | Malignant neoplasm of kidney | C04;C13;C12 | Neoplasms; Female Urogenital Diseases and Pregnancy Complications; Male Urogenital Diseases |
| C0014175 | 511 | 0.00086264 | 0.36751 | Endometriosis | C13 | Female Urogenital Diseases and Pregnancy Complications |
| C0019196 | 629 | 0.00097739 | 0.36393 | Hepatitis C | C06;C01 | Digestive System Diseases; Infections |
| C2711227 | 424 | 0.0010794 | 0.36105 | Steatohepatitis | C06 | Digestive System Diseases |
| C0027697 | 108 | 0.0012002 | 0.35795 | Nephritis | C13;C12 | Female Urogenital Diseases and Pregnancy Complications; Male Urogenital Diseases |
| C0023493 | 426 | 0.0017368 | 0.34689 | Adult T-Cell Lymphoma/Leukemia | C04;C20;C15 | Neoplasms; Immune System Diseases; Hemic and Lymphatic Diseases |
| C0001815 | 147 | 0.0017583 | 0.34651 | Primary Myelofibrosis | C15 | Hemic and Lymphatic Diseases |

| | | | | | | |
|---|---|---|---|---|---|---|
| C0036421 | 461 | 0.0017645 | 0.3464 | Systemic Scleroderma | C17 | Skin and Connective Tissue Diseases |
| C0155626 | 217 | 0.0018882 | 0.34433 | Acute myocardial infarction | C23;C14 | "Pathological Conditions, Signs and Symptoms; Cardiovascular Diseases" |
| C0677607 | 154 | 0.001936 | 0.34356 | Hashimoto Disease | C19 | Endocrine System Diseases |
| C0302592 | 907 | 0.0019578 | 0.34321 | Cervix carcinoma | C04;C13 | Neoplasms; Female Urogenital Diseases and Pregnancy Complications |
| C0024198 | 92 | 0.0019745 | 0.34295 | Lyme Disease | C01 | Infections |
| C0153676 | 580 | 0.0021391 | 0.34047 | Secondary malignant neoplasm of lung | C04;C08 | Neoplasms; Respiratory Tract Diseases |
| C0020443 | 198 | 0.0022024 | 0.33956 | Hypercholesterolemia | C18 | Nutritional and Metabolic Diseases |
| C0007222 | 657 | 0.0022104 | 0.33945 | Cardiovascular Diseases | C14 | Cardiovascular Diseases |
| C0948089 | 181 | 0.0029483 | 0.33032 | Acute Coronary Syndrome | C14 | Cardiovascular Diseases |
| C0524910 | 317 | 0.0033761 | 0.32593 | "Hepatitis C, Chronic" | C06;C01 | Digestive System Diseases; Infections |
| C0042373 | 340 | 0.0039602 | 0.32068 | Vascular Diseases | C14 | Cardiovascular Diseases |
| C0851162 | 1 | 0.0044806 | 0.31655 | Infections of musculoskeletal system | C23;C01;C05 | "Pathological Conditions, Signs and Symptoms; Infections; Musculoskeletal Diseases" |
| C0032463 | 170 | 0.0048689 | 0.31374 | Polycythemia Vera | C04;C15 | Neoplasms; Hemic and Lymphatic Diseases |

| | | | | | | |
|---|---|---|---|---|---|---|
| C0030305 | 117 | 0.0052153 | 0.31139 | Pancreatitis | C06 | Digestive System Diseases |
| C0020456 | 416 | 0.0053691 | 0.3104 | Hyperglycemia | C18 | Nutritional and Metabolic Diseases |
| C0948008 | 351 | 0.0054805 | 0.30969 | Ischemic stroke | C10;C14 | Nervous System Diseases; Cardiovascular Diseases |
| C0036220 | 279 | 0.0065327 | 0.30358 | Kaposi Sarcoma | C04;C01 | Neoplasms; Infections |
| C0699791 | 1879 | 0.0083652 | 0.29476 | Stomach Carcinoma | C06;C04 | Digestive System Diseases; Neoplasms |
| C0038454 | 589 | 0.0084979 | 0.29419 | Cerebrovascular accident | C10;C14 | Nervous System Diseases; Cardiovascular Diseases |
| C0878544 | 330 | 0.010721 | 0.28563 | Cardiomyopathies | C14 | Cardiovascular Diseases |
| C0020459 | 238 | 0.010998 | 0.28468 | Hyperinsulinism | C18 | Nutritional and Metabolic Diseases |
| C3695127 | 2 | 0.012895 | 0.27865 | Astrocytoma of brain | C04;C10 | Neoplasms; Nervous System Diseases |
| C0585442 | 819 | 0.015445 | 0.27166 | Osteosarcoma of bone | C04 | Neoplasms |
| C3887641 | 12 | 0.016258 | 0.26964 | Recurrent hepatitis | C06 | Digestive System Diseases |
| C0020473 | 158 | 0.018798 | 0.26384 | Hyperlipidemia | C18 | Nutritional and Metabolic Diseases |
| C0524620 | 461 | 0.019872 | 0.2616 | Metabolic Syndrome X | C18 | Nutritional and Metabolic Diseases |
| C0006826 | 1061 | 0.02137 | 0.25863 | Malignant Neoplasms | C04 | Neoplasms |
| C0025202 | 1930 | 0.024963 | 0.25217 | melanoma | C04 | Neoplasms |
| C0149925 | 531 | 0.026365 | 0.24986 | Small cell carcinoma of lung | C04;C08 | Neoplasms; Respiratory Tract Diseases |

| | | | | | | |
|---|---|---|---|---|---|---|
| C2939419 | 93 | 0.028499 | 0.24655 | Secondary Neoplasm | C23;C04 | "Pathological Conditions, Signs and Symptoms; Neoplasms" |
| C0035335 | 464 | 0.029609 | 0.2449 | Retinoblastoma | C04;C11 | Neoplasms; Eye Diseases |
| C0025517 | 367 | 0.033048 | 0.24012 | Metabolic Diseases | C18 | Nutritional and Metabolic Diseases |
| C0002170 | 108 | 0.033234 | 0.23987 | Alopecia | C23;C17 | "Pathological Conditions, Signs and Symptoms; Skin and Connective Tissue Diseases" |
| C0494165 | 420 | 0.033911 | 0.23899 | Secondary malignant neoplasm of liver | C06;C04 | Digestive System Diseases; Neoplasms |
| C0242339 | 174 | 0.034223 | 0.23858 | Dyslipidemias | C18 | Nutritional and Metabolic Diseases |
| C0027051 | 654 | 0.03449 | 0.23824 | Myocardial Infarction | C23;C14 | "Pathological Conditions, Signs and Symptoms; Cardiovascular Diseases" |
| C2239176 | 2810 | 0.048498 | 0.22273 | Liver carcinoma | C06;C04 | Digestive System Diseases; Neoplasms |
| C0699885 | 976 | 0.04937 | 0.22189 | Carcinoma of bladder | C04;C13;C12 | Neoplasms; Female Urogenital Diseases and Pregnancy Complications; Male Urogenital Diseases |
| C0017638 | 1748 | 0.054114 | 0.21755 | Glioma | C04 | Neoplasms |
| C0007137 | 1425 | 0.062044 | 0.21093 | Squamous cell carcinoma | C04 | Neoplasms |
| C0010054 | 645 | 0.078594 | 0.19908 | Coronary Arteriosclerosis | C14 | Cardiovascular Diseases |

| | | | | | | |
|---|---|---|---|---|---|---|
| C0271650 | 261 | 0.085441 | 0.19476 | Impaired glucose tolerance | C18 | Nutritional and Metabolic Diseases |
| C1306459 | 847 | 0.096692 | 0.18821 | Primary malignant neoplasm | C04 | Neoplasms |
| C0011849 | 1243 | 0.10795 | 0.18224 | Diabetes Mellitus | C18;C19 | Nutritional and Metabolic Diseases; Endocrine System Diseases |
| C1140680 | 1668 | 0.1138 | 0.17933 | Malignant neoplasm of ovary | C04;C13;C19 | Neoplasms; Female Urogenital Diseases and Pregnancy Complications; Endocrine System Diseases |
| C0007131 | 1784 | 0.11705 | 0.17776 | Non-Small Cell Lung Carcinoma | C04;C08 | Neoplasms; Respiratory Tract Diseases |
| C0027819 | 1445 | 0.12619 | 0.17352 | Neuroblastoma | C04 | Neoplasms |
| C0699790 | 1771 | 0.12619 | 0.17352 | Colon Carcinoma | C06;C04 | Digestive System Diseases; Neoplasms |
| C0242379 | 1946 | 0.14719 | 0.1646 | Malignant neoplasm of lung | C04;C08 | Neoplasms; Respiratory Tract Diseases |
| C0043167 | 10 | 0.14894 | -0.16389 | Pertussis | C01;C08 | Infections; Respiratory Tract Diseases |
| C0018802 | 629 | 0.15792 | 0.1604 | Congestive heart failure | C14 | Cardiovascular Diseases |
| C0011860 | 1232 | 0.28004 | 0.12304 | Diabetes Mellitus, Non-Insulin-Dependent | C18;C19 | Nutritional and Metabolic Diseases; Endocrine System Diseases |
| C0235974 | 1552 | 0.3264 | 0.11185 | Pancreatic carcinoma | C06;C04;C19 | Digestive System Diseases; Neoplasms; |

| | | | | | | |
|---|---|---|---|---|---|---|
| | | | | | | Endocrine System Diseases |
| C0024121 | 598 | 0.35588 | 0.10526 | Lung Neoplasms | C04;C08 | Neoplasms; Respiratory Tract Diseases |
| C0001418 | 1298 | 0.35951 | 0.10447 | Adenocarcinoma | C04 | Neoplasms |
| C1458155 | 1506 | 0.41849 | 0.092302 | Mammary Neoplasms | C04;C17 | Neoplasms; Skin and Connective Tissue Diseases |
| C0686619 | 1039 | 0.43439 | 0.089195 | Secondary malignant neoplasm of lymph node | C23;C04 | "Pathological Conditions, Signs and Symptoms; Neoplasms" |
| C0678222 | 3876 | 0.5947 | 0.060772 | Breast Carcinoma | C04;C17 | Neoplasms; Skin and Connective Tissue Diseases |
| C0006142 | 3779 | 0.62734 | 0.055461 | Malignant neoplasm of breast | C04;C17 | Neoplasms; Skin and Connective Tissue Diseases |
| C0001430 | 828 | 0.79925 | -0.02907 | Adenoma | C04 | Neoplasms |
| C0020538 | 876 | 0.8853 | -0.016492 | Hypertensive disease | C14 | Cardiovascular Diseases |
| C0376358 | 2468 | 0.98317 | -0.0024114 | Malignant neoplasm of prostate | C04;C12 | Neoplasms; Male Urogenital Diseases |

Table S4.  The 171 diseases classified into three clusters based on their cytokine profiles

| Concept | Disease/Symptom | Class | Class Name |
|---|---|---|---|
| Cluster 1-2 (blue and cyan) | | | |
| C0014175 | Endometriosis | C13 | Female Urogenital Diseases and Pregnancy Complications |
| C0010054 | Coronary Arteriosclerosis | C14 | Cardiovascular Diseases |
| C0042373 | Vascular Diseases | C14 | Cardiovascular Diseases |
| C0948008 | Ischemic stroke | C10;C14 | Nervous System Diseases; Cardiovascular Diseases |
| C0027051 | Myocardial Infarction | C23;C14 | Pathological Conditions, Signs and Symptoms; Cardiovascular Diseases |
| C0038454 | Cerebrovascular accident | C10;C14 | Nervous System Diseases; Cardiovascular Diseases |
| C0007222 | Cardiovascular Diseases | C14 | Cardiovascular Diseases |
| C0018802 | Congestive heart failure | C14 | Cardiovascular Diseases |
| C0524620 | Metabolic Syndrome X | C18 | Nutritional and Metabolic Diseases |
| C0020456 | Hyperglycemia | C18 | Nutritional and Metabolic Diseases |
| C0020459 | Hyperinsulinism | C18 | Nutritional and Metabolic Diseases |
| C0020443 | Hypercholesterolemia | C18 | Nutritional and Metabolic Diseases |
| C0240066 | Iron deficiency | C18 | Nutritional and Metabolic Diseases |
| C0008354 | Cholera | C01 | Infections |
| C0153690 | Secondary malignant neoplasm of bone | C23;C04;C05 | Pathological Conditions, Signs and Symptoms; Neoplasms; Musculoskeletal Diseases |
| C0153676 | Secondary malignant neoplasm of lung | C04;C08 | Neoplasms; Respiratory Tract Diseases |
| C2939419 | Secondary Neoplasm | C23;C04 | Pathological Conditions, Signs and Symptoms; Neoplasms |
| C0494165 | Secondary malignant neoplasm of liver | C06;C04 | Digestive System Diseases; Neoplasms |
| C0023440 | Acute Erythroblastic Leukemia | C04;C15 | Neoplasms; Hemic and Lymphatic Diseases |
| C0271650 | Impaired glucose tolerance | C18 | Nutritional and Metabolic Diseases |

| | | | |
|---|---|---|---|
| C0020538 | Hypertensive disease | C14 | Cardiovascular Diseases |
| C0020473 | Hyperlipidemia | C18 | Nutritional and Metabolic Diseases |
| C0242339 | Dyslipidemias | C18 | Nutritional and Metabolic Diseases |
| C0018799 | Heart Diseases | C14 | Cardiovascular Diseases |
| C0035335 | Retinoblastoma | C04;C11 | Neoplasms; Eye Diseases |
| C0027819 | Neuroblastoma | C04 | Neoplasms |
| C1458155 | Mammary Neoplasms | C04;C17 | Neoplasms; Skin and Connective Tissue Diseases |
| C1140680 | Malignant neoplasm of ovary | C04;C13;C19 | Neoplasms; Female Urogenital Diseases and Pregnancy Complications; Endocrine System Diseases |
| C0017638 | Glioma | C04 | Neoplasms |
| C0007137 | Squamous cell carcinoma | C04 | Neoplasms |
| C0007131 | Non-Small Cell Lung Carcinoma | C04;C08 | Neoplasms; Respiratory Tract Diseases |
| C0699790 | Colon Carcinoma | C06;C04 | Digestive System Diseases; Neoplasms |
| C0699791 | Stomach Carcinoma | C06;C04 | Digestive System Diseases; Neoplasms |
| C0001418 | Adenocarcinoma | C04 | Neoplasms |
| C0686619 | Secondary malignant neoplasm of lymph node | C23;C04 | Pathological Conditions, Signs and Symptoms; Neoplasms |
| C0699885 | Carcinoma of bladder | C04;C13;C12 | Neoplasms; Female Urogenital Diseases and Pregnancy Complications; Male Urogenital Diseases |
| C0235974 | Pancreatic carcinoma | C06;C04;C19 | Digestive System Diseases; Neoplasms; Endocrine System Diseases |
| C0302592 | Cervix carcinoma | C04;C13 | Neoplasms; Female Urogenital Diseases and Pregnancy Complications |
| C0025202 | melanoma | C04 | Neoplasms |
| C1306459 | Primary malignant neoplasm | C04 | Neoplasms |
| C0006826 | Malignant Neoplasms | C04 | Neoplasms |
| C0740457 | Malignant neoplasm of kidney | C04;C13;C12 | Neoplasms; Female Urogenital Diseases and Pregnancy |

| | | | |
|---|---|---|---|
| | | | Complications; Male Urogenital Diseases |
| C0585442 | Osteosarcoma of bone | C04 | Neoplasms |
| C0149925 | Small cell carcinoma of lung | C04;C08 | Neoplasms; Respiratory Tract Diseases |
| C0024121 | Lung Neoplasms | C04;C08 | Neoplasms; Respiratory Tract Diseases |
| C2711227 | Steatohepatitis | C06 | Digestive System Diseases |
| C0011849 | Diabetes Mellitus | C18;C19 | Nutritional and Metabolic Diseases; Endocrine System Diseases |
| C0011860 | Diabetes Mellitus, Non-Insulin-Dependent | C18;C19 | Nutritional and Metabolic Diseases; Endocrine System Diseases |
| C0025517 | Metabolic Diseases | C18 | Nutritional and Metabolic Diseases |
| C0001430 | Adenoma | C04 | Neoplasms |
| C0878544 | Cardiomyopathies | C14 | Cardiovascular Diseases |
| C0242379 | Malignant neoplasm of lung | C04;C08 | Neoplasms; Respiratory Tract Diseases |
| C0376358 | Malignant neoplasm of prostate | C04;C12 | Neoplasms; Male Urogenital Diseases |
| C2239176 | Liver carcinoma | C06;C04 | Digestive System Diseases; Neoplasms |
| C0006142 | Malignant neoplasm of breast | C04;C17 | Neoplasms; Skin and Connective Tissue Diseases |
| C0678222 | Breast Carcinoma | C04;C17 | Neoplasms; Skin and Connective Tissue Diseases |
| Cluster 3 (green) | | | |
| C0036421 | Systemic Scleroderma | C17 | Skin and Connective Tissue Diseases |
| C0035243 | Respiratory Tract Infections | C01;C08 | Infections; Respiratory Tract Diseases |
| C0036690 | Septicemia | C23;C01 | Pathological Conditions, Signs and Symptoms; Infections |
| C0243026 | Sepsis | C23;C01 | Pathological Conditions, Signs and Symptoms; Infections |
| C0002940 | Aneurysm | C14 | Cardiovascular Diseases |
| C0948089 | Acute Coronary Syndrome | C14 | Cardiovascular Diseases |
| C0025289 | Meningitis | C10 | Nervous System Diseases |

| | | | |
|---|---|---|---|
| C0023281 | Leishmaniasis | C01;C17 | Infections; Skin and Connective Tissue Diseases |
| C0003872 | Arthritis, Psoriatic | C17;C05 | Skin and Connective Tissue Diseases; Musculoskeletal Diseases |
| C1290884 | Inflammatory disorder | C23 | Pathological Conditions, Signs and Symptoms |
| C0023343 | Leprosy | C01 | Infections |
| C0042164 | Uveitis | C11 | Eye Diseases |
| C0011603 | Dermatitis | C17 | Skin and Connective Tissue Diseases |
| C0876973 | Infectious Lung Disorder | C01;C08 | Infections; Respiratory Tract Diseases |
| C0017152 | Gastritis | C06 | Digestive System Diseases |
| C0017658 | Glomerulonephritis | C13;C12 | Female Urogenital Diseases and Pregnancy Complications; Male Urogenital Diseases |
| C0031099 | Periodontitis | C07 | Stomatognathic Diseases |
| C0266929 | Chronic Periodontitis | C07 | Stomatognathic Diseases |
| C0039103 | Synovitis | C05 | Musculoskeletal Diseases |
| C0031154 | Peritonitis | C06;C01 | Digestive System Diseases; Infections |
| C0027697 | Nephritis | C13;C12 | Female Urogenital Diseases and Pregnancy Complications; Male Urogenital Diseases |
| C0024143 | Lupus Nephritis | C13;C17;C12;C20 | Female Urogenital Diseases and Pregnancy Complications; Skin and Connective Tissue Diseases; Male Urogenital Diseases; Immune System Diseases |
| C1290886 | Chronic inflammatory disorder | C23 | Pathological Conditions, Signs and Symptoms |
| C0029118 | Opportunistic Infections | C01 | Infections |
| C0242966 | Systemic Inflammatory Response Syndrome | C23 | Pathological Conditions, Signs and Symptoms |
| C0001339 | Acute pancreatitis | C06 | Digestive System Diseases |
| C0026946 | Mycoses | C01 | Infections |
| C0011615 | Dermatitis, Atopic | C16;C17;C20 | Congenital, Hereditary, and Neonatal Diseases and Abnormalities; Skin and |

| | | | |
|---|---|---|---|
| | | | Connective Tissue Diseases; Immune System Diseases |
| C0033860 | Psoriasis | C17 | Skin and Connective Tissue Diseases |
| C0524910 | Hepatitis C, Chronic | C06;C01 | Digestive System Diseases; Infections |
| C0018213 | Graves Disease | C11;C20;C19 | Eye Diseases; Immune System Diseases; Endocrine System Diseases |
| C0041327 | Tuberculosis, Pulmonary | C01;C08 | Infections; Respiratory Tract Diseases |
| C0035235 | Respiratory Syncytial Virus Infections | C01 | Infections |
| C1306759 | Eosinophilic disorder | C15 | Hemic and Lymphatic Diseases |
| C0027059 | Myocarditis | C14 | Cardiovascular Diseases |
| C2607914 | Allergic rhinitis (disorder) | C08;C20;C09 | Respiratory Tract Diseases; Immune System Diseases; Otorhinolaryngologic Diseases |
| C0155877 | Allergic asthma | C08;C20 | Respiratory Tract Diseases; Immune System Diseases |
| C0042384 | Vasculitis | C14 | Cardiovascular Diseases |
| C3714636 | Pneumonitis | C01;C08 | Infections; Respiratory Tract Diseases |
| C0001175 | Acquired Immunodeficiency Syndrome | C01;C20 | Infections; Immune System Diseases |
| C0036220 | Kaposi Sarcoma | C04;C01 | Neoplasms; Infections |
| C0026948 | Mycosis Fungoides | C04;C20;C15 | Neoplasms; Immune System Diseases; Hemic and Lymphatic Diseases |
| C0014038 | Encephalitis | C10 | Nervous System Diseases |
| C0036202 | Sarcoidosis | C15 | Hemic and Lymphatic Diseases |
| C0149516 | Chronic sinusitis | C01;C08;C09 | Infections; Respiratory Tract Diseases; Otorhinolaryngologic Diseases |
| C3812396 | Chronic idiopathic pulmonary fibrosis | C08 | Respiratory Tract Diseases |
| C0242497 | Intestinal schistosomiasis | C01 | Infections |
| C3695127 | Astrocytoma of brain | C04;C10 | Neoplasms; Nervous System Diseases |

| | | | |
|---|---|---|---|
| C0155626 | Acute myocardial infarction | C23;C14 | Pathological Conditions, Signs and Symptoms; Cardiovascular Diseases |
| Cluster 4-5 (red and orange) | | | |
| C0001486 | Adenovirus Infections | C01 | Infections |
| C0006118 | Brain Neoplasms | C04;C10 | Neoplasms; Nervous System Diseases |
| C0023487 | Acute Promyelocytic Leukemia | C04 | Neoplasms |
| C0023473 | Myeloid Leukemia, Chronic | C04;C15 | Neoplasms; Hemic and Lymphatic Diseases |
| C0023467 | Leukemia, Myelocytic, Acute | C04 | Neoplasms |
| C0023418 | leukemia | C04 | Neoplasms |
| C0023449 | Acute lymphocytic leukemia | C04;C20;C15 | Neoplasms; Immune System Diseases; Hemic and Lymphatic Diseases |
| C0019163 | Hepatitis B | C06;C01 | Digestive System Diseases; Infections |
| C3463824 | MYELODYSPLASTIC SYNDROME | C15 | Hemic and Lymphatic Diseases |
| C0085669 | Acute leukemia | C23;C04 | Pathological Conditions, Signs and Symptoms; Neoplasms |
| C0023470 | Myeloid Leukemia | C04 | Neoplasms |
| C0376545 | Hematologic Neoplasms | C04;C15 | Neoplasms; Hemic and Lymphatic Diseases |
| C0006413 | Burkitt Lymphoma | C04;C01;C20;C15 | Neoplasms; Infections; Immune System Diseases; Hemic and Lymphatic Diseases |
| C0079731 | B-Cell Lymphomas | C04;C20;C15 | Neoplasms; Immune System Diseases; Hemic and Lymphatic Diseases |
| C0023434 | Chronic Lymphocytic Leukemia | C04;C20;C15 | Neoplasms; Immune System Diseases; Hemic and Lymphatic Diseases |
| C0024299 | Lymphoma | C04;C20;C15 | Neoplasms; Immune System Diseases; Hemic and Lymphatic Diseases |
| C0026764 | Multiple Myeloma | C04;C20;C15;C14 | Neoplasms; Immune System Diseases; Hemic and Lymphatic Diseases; Cardiovascular Diseases |

| | | | |
|---|---|---|---|
| C0024305 | Lymphoma, Non-Hodgkin | C04;C20;C15 | Neoplasms; Immune System Diseases; Hemic and Lymphatic Diseases |
| C0030305 | Pancreatitis | C06 | Digestive System Diseases |
| C0024530 | Malaria | C01 | Infections |
| C0275524 | Coinfection | C01 | Infections |
| C0010823 | Cytomegalovirus Infections | C01 | Infections |
| C0019693 | HIV Infections | C01;C20 | Infections; Immune System Diseases |
| C0009324 | Ulcerative Colitis | C06 | Digestive System Diseases |
| C0021390 | Inflammatory Bowel Diseases | C06 | Digestive System Diseases |
| C0010346 | Crohn Disease | C06 | Digestive System Diseases |
| C0026769 | Multiple Sclerosis | C20;C10 | Immune System Diseases; Nervous System Diseases |
| C0003864 | Arthritis | C05 | Musculoskeletal Diseases |
| C0036323 | Schistosomiasis | C01 | Infections |
| C0677607 | Hashimoto Disease | C19 | Endocrine System Diseases |
| C0026918 | Mycobacterium Infections | C01 | Infections |
| C0007570 | Celiac Disease | C06;C18 | Digestive System Diseases; Nutritional and Metabolic Diseases |
| C0023493 | Adult T-Cell Lymphoma/Leukemia | C04;C20;C15 | Neoplasms; Immune System Diseases; Hemic and Lymphatic Diseases |
| C0009319 | Colitis | C06 | Digestive System Diseases |
| C0019159 | Hepatitis A | C06;C01 | Digestive System Diseases; Infections |
| C0041296 | Tuberculosis | C01 | Infections |
| C0524909 | Hepatitis B, Chronic | C06;C01 | Digestive System Diseases; Infections |
| C0024198 | Lyme Disease | C01 | Infections |
| C0011311 | Dengue Fever | C01 | Infections |
| C0025007 | Measles | C01 | Infections |
| C0038013 | Ankylosing spondylitis | C05 | Musculoskeletal Diseases |
| C0024131 | Lupus Vulgaris | C01;C17 | Infections; Skin and Connective Tissue Diseases |
| C0019829 | Hodgkin Disease | C04;C20;C15 | Neoplasms; Immune System Diseases; Hemic and Lymphatic Diseases |

| | | | |
|---|---|---|---|
| C0023443 | Hairy Cell Leukemia | C04;C20;C15 | Neoplasms; Immune System Diseases; Hemic and Lymphatic Diseases |
| C0040100 | Thymoma | C04;C15 | Neoplasms; Hemic and Lymphatic Diseases |
| C0032463 | Polycythemia Vera | C04;C15 | Neoplasms; Hemic and Lymphatic Diseases |
| C0002170 | Alopecia | C23;C17 | Pathological Conditions, Signs and Symptoms; Skin and Connective Tissue Diseases |
| C1264606 | Persistent infection | C23;C01 | Pathological Conditions, Signs and Symptoms; Infections |
| C0019348 | Herpes Simplex Infections | C01;C17 | Infections; Skin and Connective Tissue Diseases |
| C0003873 | Rheumatoid Arthritis | C17;C05;C20 | Skin and Connective Tissue Diseases; Musculoskeletal Diseases; Immune System Diseases |
| C0011854 | Diabetes Mellitus, Insulin-Dependent | C18;C20;C19 | Nutritional and Metabolic Diseases; Immune System Diseases; Endocrine System Diseases |
| C0019196 | Hepatitis C | C06;C01 | Digestive System Diseases; Infections |
| C0042769 | Virus Diseases | C01 | Infections |
| C0024141 | Lupus Erythematosus, Systemic | C17;C20 | Skin and Connective Tissue Diseases; Immune System Diseases |
| C0021400 | Influenza | C01;C08 | Infections; Respiratory Tract Diseases |
| C0266999 | Vesicular Stomatitis | C01;C07;C22 | Infections; Stomatognathic Diseases; Animal Diseases |
| C0024266 | Lymphocytic Choriomeningitis | C01;C10 | Infections; Nervous System Diseases |
| C0036117 | Salmonella infections | C01 | Infections |
| C0023290 | Leishmaniasis, Visceral | C01 | Infections |
| C0004623 | Bacterial Infections | C01 | Infections |
| C0026936 | Mycoplasma Infections | C01 | Infections |
| C0151317 | Chronic infectious disease | C20 | Immune System Diseases |
| C3887641 | Recurrent hepatitis | C06 | Digestive System Diseases |

| | | | |
|---|---|---|---|
| C0001815 | Primary Myelofibrosis | C15 | Hemic and Lymphatic Diseases |
| Other (top and bottom, dark) | | | |
| C0851162 | Infections of musculoskeletal system | C23;C01;C05 | Pathological Conditions, Signs and Symptoms; Infections; Musculoskeletal Diseases |
| C0043167 | Pertussis | C01;C08 | Infections; Respiratory Tract Diseases |

Table S5. Disease-associated genes in the well-connected modules formed by pathogenesis genes, receptors, and essential cytokines identified by spectrum partition on the interactions between disease-specific cytokine networks in the context of five immune disorders: rheumatoid arthritis (RA), psoriasis (PS), ulcerative colitis (UC), Crohn's disease (CD) and systemic lupus erythematosus (SLE). Essential cytokines are marked with "E"; cytokine receptors are marked with "R"; other genes are marked with "D".

| rheumatoid arthritis | | psoriasis | systemic lupus erythematosus | | ulcerative colitis | Crohn's disease | |
|---|---|---|---|---|---|---|---|
| TNFRSF25 R<br>TRAF1 D<br>FASLG D<br>TNFRSF9 R<br>IL1A E<br>MAP3K5 D<br>TNFRSF1A R<br>MALT1 D<br>TLR4 D<br>TLR5 D<br>IL1RN R<br>IRAK1 D<br>S100A8 D<br>TLR1 D | IL7 E<br>STAT3 D<br>TYK2 D<br>IL4R R<br>IL9 E<br>OSMR D<br>JAK1 D<br>IL23A E<br>IL2 E<br>STAT5A D<br>STAT5B D<br>IFNGR1 R<br>STAT4 D<br>IL6R R<br>IL15 E<br>IFNGR2 R<br>IL2RA R<br>LIF E<br>CD70 E<br>IL2RG R | FASLG D<br>TNFRSF9 R<br>TNFSF15 E<br>TNFRSF17 R<br>TNFRSF10B R<br>TNFRSF13B R<br>TNFRSF6B R<br>TNFSF14 E<br>PSMD7 D<br>REL D<br>LTBR D<br>TNFRSF13C R<br>TNFSF13B E<br>TNFSF11 E | TRAF1 D<br>FASLG D<br>IL1A E<br>TNFRSF1A R<br>TLR4 D<br>IL1RN R<br>TLR5 D<br>IRAK1 D<br>S100A8 D<br>NLRP3 D<br>TIRAP D<br>MYD88 D<br>TNF E<br>TLR2 D<br>TLR9 D<br>IL18R1 R<br>IRAK4 D<br>NFKBIA D<br>IL1R1 R<br>IL33 E<br>IL1B E<br>SIGIRR D<br>IL18 E<br>TNFSF4 E<br>TNFRSF4 R<br>IL12RB1 R<br>FOXP3 D<br>IL23R R<br>IL17A E<br>TBX21 D<br>IL13 E | IL2RA R<br>IL4R R<br>TYK2 D<br>IFNL1 E<br>IL26 E<br>IL6R R<br>OSMR D<br>IL21 E<br>IL27 E<br>LIF E<br>IFNGR1 R<br>IFNGR2 R<br>STAT4 D<br>IL24 E<br>IL31 E<br>PRL D<br>IL21R R<br>IFNL2 E<br>STAT1 D<br>IL20 E<br>IFNLR1 R<br>IKZF3 D | TNFRSF9 R<br>TNFSF15 E<br>TNFRSF17 R<br>CFLAR D<br>TNFRSF6B R<br>TNFSF14 E<br>BIRC2 D<br>BIRC3 D<br>NFKB2 D<br>USP14 D<br>REL D<br>PSMG1 D<br>EGLN3 D<br>TNFSF13B E<br>TNFRSF11A R<br>TNFSF11 E | TNFRSF9 R<br>IL1A E<br>MAP3K1 D<br>TNFRSF1A R<br>TNFSF4 E<br>TLR4 D<br>TLR5 D<br>IRAK1 D<br>TLR1 D<br>IL1RL1 R<br>IRAK3 D<br>NLRP3 D<br>IL1RN R<br>IL18RAP R<br>TLR2 D | IL19 E<br>IL15 E<br>IL27 E<br>IL11 E<br>IL12RB2 R<br>IL7 E<br>IL5 E<br>IL12B E<br>IFNL1 E<br>IL23A E<br>IL4R R<br>IL9 E<br>TYK2 D<br>STAT3 D<br>GHR D<br>STAT5B D<br>STAT5A D<br>STAT4 D<br>LIF E<br>IFNGR2 R<br>IL2RA R<br>IL15RA R |

| | | | | | | | |
|---|---|---|---|---|---|---|---|
| NLRP3<br>D<br>TIRAP<br>D<br>MYD88<br>D<br>TNF E<br>TLR2<br>D<br>TLR9<br>D<br>IL18RAP R<br>IL18R1 R<br>IL33 E<br>NFKBIA<br>D<br>IL1R1 R<br>SIGIRR<br>D<br>IL18 E<br>IL1B E<br>TNFSF4 E<br>TSLP<br>D<br>FOXP3<br>D<br>IL23R R<br>STAT6<br>D<br>IL17A E | STAT1<br>D<br>IL31 E<br>IL2RB R<br>IL3 E<br>IFNL2 E<br>IL21R R<br>GH1<br>D<br>IFNLR1 R<br>IL27RA R<br>CSF2RA R<br>IL5RA R<br>IL9R R<br>IFNA6 E<br>IFNA2 E<br>PRL<br>D<br>IKZF3<br>D<br>BCL6<br>D<br>SH2B3<br>D<br>SELE<br>D<br>IFNA1 E<br>CTLA4<br>D<br>CR2<br>D<br>PTPN2<br>D | | IL15 E<br>IL19 E<br>IL10RA R<br>IL10 E<br>IL2 E<br>IL4 E<br>IL22 E<br>IL23A E<br>IL12B E<br>IL12A E<br>IL10RB R<br>IL7 E<br>IL9 E<br>IL5 E<br>IL11 E<br>IFNG E<br>IL3 E<br>JAK1 D<br>STAT3 D | CD70 E<br>SH2B3<br>D<br>IFNAR2 R<br>CR2<br>D<br>STAT2<br>D<br>SELE<br>D<br>IFNA2 E<br>IFNA6 E<br>CTLA4<br>D<br>IFNA1 E<br>IFNK E<br>IRF9<br>D<br>IRF7<br>D<br>HLX<br>D<br>IFNB1 E<br>IFIT1<br>D<br>IFI44<br>D<br>IRF1<br>D<br>IRF2<br>D | | TLR9<br>D<br>TNF E<br>IL33 E<br>NFKBIA<br>D<br>IL18 E<br>IL1B E<br>FOXP3<br>D<br>TSLP<br>D<br>STAT6<br>D<br>IL23R R<br>IL10RA R<br>IL17A E<br>IL10 E<br>IL13 E<br>IL10RB R<br>IL12A E<br>IL4 E<br>IL21 E<br>IFNG F<br>IL24 E<br>IL20 E<br>IL26 E<br>IL22 E | IL31 E<br>IFNL2 F<br>IL2 E<br>IL3 E<br>CD70 E<br>GH1<br>D<br>JAK2<br>D<br>CSF2RA R<br>IFNA2 E<br>IFNA6 E<br>IFNA1 E<br>PTPN2<br>D<br>IFNL3 E<br>CTLA4<br>D<br>INPP5D<br>D |

| | | | | |
|---|---|---|---|---|
| TBX21<br><br>D<br>IL13 E<br>IL10 E<br>IL10RA R<br>IL10RB R<br>IL4 E<br>IL12A E<br>IFNG E<br>IL21 E<br>IL24 E<br>IL19 E<br>IL20 E<br>IL22 E<br>IL26 E<br>IL11 E<br>IL12B E<br>IFNL1 E<br>IL27 E<br>IL5 E | INPP5D<br><br>D<br>IFI44<br><br>D<br>IFNL3 E<br>CLEC7A<br><br>D<br>TNFSF9<br>E<br>MAP4K3<br><br>D<br>PDCD5<br><br>D<br>SPATA2<br><br>D<br>TNIP1<br><br>D<br>DDAH1<br><br>D | | | IRF5<br><br>D<br>IFNL3 E<br>CLEC7A<br><br>D<br>TNFSF9<br>E<br>MAP4K3<br><br>D<br>SPATA2<br><br>D<br>TNIP1<br><br>D<br>ZC3H12<br>A<br><br>D |
